# inVAE: Conditionally invariant representation learning for generating multivariate single-cell reference maps

**DOI:** 10.1101/2024.12.06.627196

**Authors:** Hananeh Aliee, Ferdinand Kapl, Duy Pham, Batuhan Cakir, Takahiro Jimba, James Cranley, Sarah A. Teichmann, Kerstin B. Meyer, Roser Vento-Tormo, Fabian J. Theis

## Abstract

Single-cell data is driving new insights into the spatiotemporal dynamics of cells and individual disease susceptibility. However, accurately identifying cell states across diverse cohorts remains challenging, as both biological variation and technical biases cause distributional shifts in the data. Separating these effects is crucial for capturing cellular heterogeneity and ensuring interpretability. To address this, we developed *inVAE*, a conditionally invariant deep generative model based on variational autoencoders. inVAE models the latent space as a combination of invariant variables, encoding true biological signals, and spurious variables, capturing technical biases. By conditioning the prior distribution of cells on biological covariates, such as disease variants, inVAE identifies high-resolution cell states in the invariant representation. Enforcing independence between the two representations disentangles biological signals from noise, enabling a more interpretable and generalizable model with a causal semantic. inVAE outperformed existing methods across four human cellular atlases of the human heart and lung, while uncovering novel cell states. It precisely stratified cell atlas donors based on the genetic impact of pathogenic variants, and excelled in predicting cell types and disease in unseen data, proving its generalizability as a reference model for label transfer. Furthermore, inVAE accurately identified temporal cell states and trajectories from developmental datasets, and captured spatial cell states in a spatially-resolved atlas. In summary, inVAE provides a powerful method for integrating multivariate single-cell transcriptomics data. By leveraging prior knowledge such as metadata, it effectively accounts for biological variation and improves latent space interpretability by disentangling biological and technical sources of variation. These capabilities enable deeper insights into cellular heterogeneity and its role in disease progression.

## Main

Single-cell transcriptomics datasets are increasing in number and complexity, including diverse cell states derived from multiple diseases, developmental stages and spatial locations^1,2^. Integrating this data is essential for capturing the full spectrum of cellular diversity, particularly in human studies, where limited sample availability and finite biological material restrict the experimental conditions that can be examined within a single study or lab. By building integrated cellular atlases from multiple studies, we can better identify inter-individual variation and the molecular characteristics unique to specific conditions^3,4^. The resulting integrated atlases have led to discoveries of previously unknown cell types^5–7^, enhanced patient stratification^8–10^, enabled robust analyses of differentially expressed genes (DEGs) across donors^11^, identified reproducible marker genes^12^, and facilitated comparisons between in vivo and in vitro models^13,14^ and between species^15–17^.

Building comprehensive atlases involves data collection under varying experimental protocols and technologies, which introduces technical biases, commonly known as batch effects. Batch effects make the separation of noise from key biological signals exceptionally challenging. Numerous computational methods have been developed to address this issue, allowing us to capture cellular phenotypes more accurately while minimising technical variation^18–23^. Among them, machine learning-based data integration methods have gained attention both due to their performance and because they enable a wide range of tasks. These tasks include mapping high-dimensional data to lower-dimensional space (representation learning), performing differential expression testing, reducing noise in data (denoising), mapping new datasets to the same low-dimensional space of a reference atlas (reference mapping), and performing automated cell-type annotation (label transfer)^19,24–28^.

The existing integration methods often remove the impact of spurious correlation, like batch effects, assuming that the underlying distribution of cells—once isolated from these confounding effects—remains consistent across datasets^19^. This assumption is based on the idea that invariant causal mechanisms are present across datasets, enabling the model to generalise to new data. However, in biological contexts, causal mechanisms can change over time or across different experimental conditions, which can invalidate the assumption of uniform distribution^29^. For instance, the causal relationship between a gene and a phenotype might depend on disease variants that vary across donors. Therefore, these methods often fail to capture critical biological variations unique to individual donors or specific conditions.

Given this assumption underlying the existing integration methods, a common approach to identify disease-specific cell states is to build atlases using similar datasets (e.g., all healthy donors) and apply reference mapping algorithms to project more distinct samples (e.g., diseased samples) onto the reference atlases^4,30^. However, the effectiveness of this approach is limited by the models’ capacity to serve as comprehensive references and the performance of the reference-mapping algorithms. This underscores the need for more generalizable integration methods that can build reference models capable of spanning a broader range of biological conditions and supporting more robust reference mapping.

Here, we developed a *conditionally* invariant deep generative model based on variational autoencoders (VAEs)^31^, named inVAE, to integrate diverse single-cell transcriptomics datasets. To accurately dissect the sources of variation across cells, inVAE infers two sets of latent variables, invariant, capturing true biological signals, and spurious, representing noise or technical biases. inVAE further incorporates prior knowledge, such as metadata, into the inference process, learning prior distributions of cells specific to each biological condition, like disease variants, developmental stage, or anatomical region. This is crucial to discover high-resolution cell states and generate representative reference atlases from multivariate data which accounts for both shared and unique molecular signatures across individuals. Moreover, inVAE has built-in predictors that transfer labels to new datasets and, crucially, identifies novel cell states when the cells in a new dataset undergo domain shifts, such as those caused by previously unseen diseases.

We applied inVAE to construct four reference atlases in the heart and the lung, providing valuable resources for future research. Our results show that inVAE effectively stratified human heart cell atlas donors by the genetic impact of pathogenic variants and excelled in predicting cell types and disease in unseen datasets, demonstrating its generalizability for label transfer. Additionally, it accurately identified temporal cell states and derived realistic lineage trajectories from developmental human lung datasets, as well as capturing spatial cell states in a spatially-resolved adult human lung atlas. Recognizing that experimental validation is not always feasible, we also provide a comprehensive set of computational methods to increase confidence in cell state identification.

## Results

### Conditionally invariant representation learning for disentangling biological and technical variation

A schematic overview of the proposed invariant variational autoencoder (inVAE) model is presented in Figure 1 (see also Supplementary Note, Box 1). The model inputs consist of the gene expression vector *X* and its associated domain *U*, indicating the conditions under which the sample was collected (i.e. metadata). The domain *U* = (*B*, *T*) encompasses two categories of observed variables: biologically relevant (*B*) and technical (*T*) variables. The biological variables *B* may include disease variants, developmental stages, anatomical tissue locations, perturbations, or other concurrently observed factors that could drive cellular changes (Figure 1c). In contrast, the technical variables *T* capture unwanted variability or biases within the data, such as batch effects or confounding factors like sex or age, depending on the experimental design.

**Figure 1:**
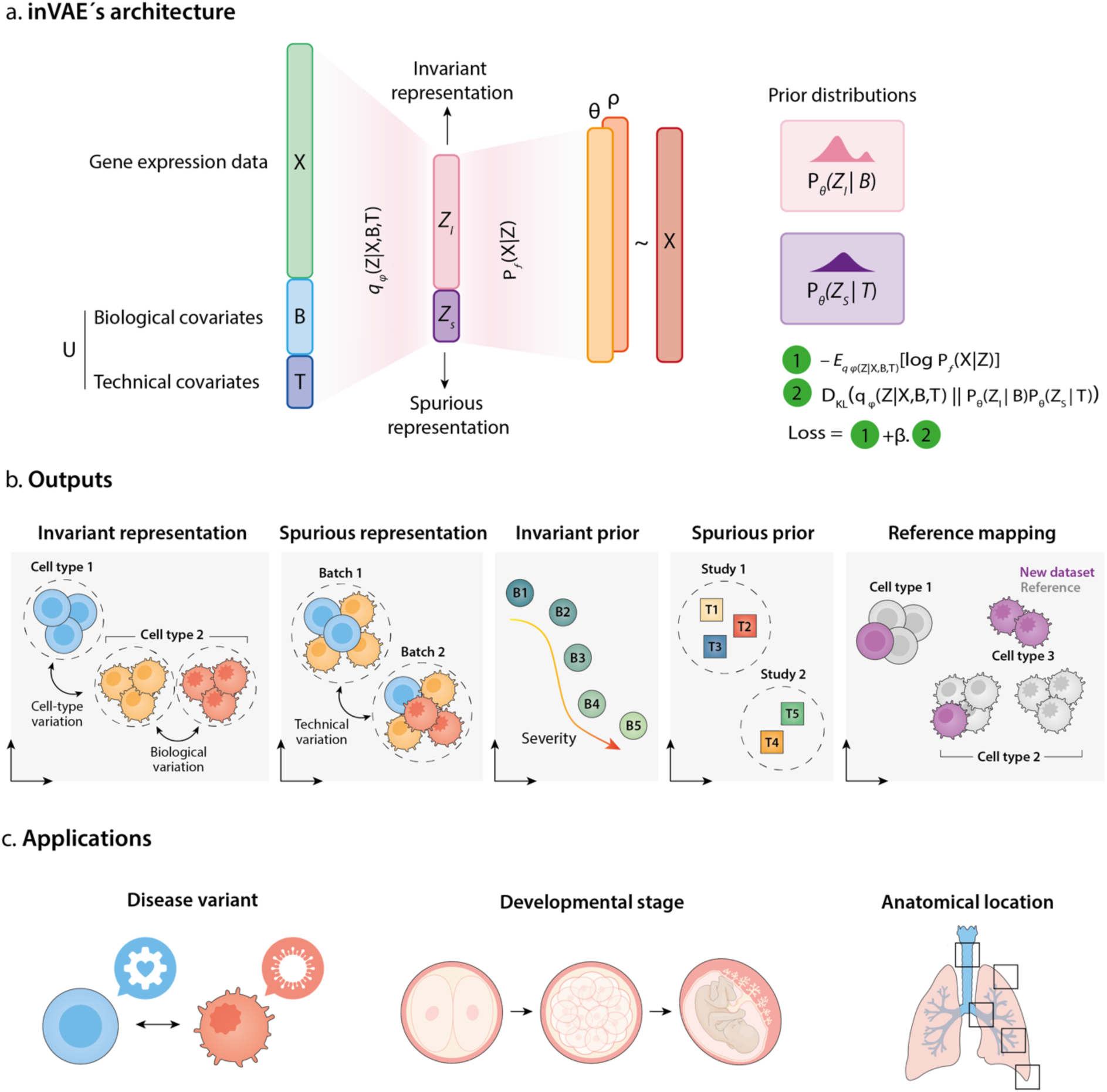
Schematic overview of the inVAE framework. **a**, The inVAE model incorporates both technical (T) and biological (B) covariates, using a nonlinear encoder to learn both invariant and spurious latent representations from high-dimensional data (X). Conditional prior distributions of cells are inferred via separate encoders, while a nonlinear decoder estimates the parameters of negative binomial distributions. The model’s loss function consists of two terms: (1) the negative log-likelihood, assessing the difference between the reconstructed and original count matrices, and (2) the KL divergence, measuring the divergence between the posterior *q_ϕ_* and prior *p_θ_* distributions. **b,** inVAE generates a range of outputs, including invariant and spurious representations of the data, which can be leveraged for downstream analysis such as data visualisation.

#### Box 1: Machine learning terminology

##### Generative model

A generative model is a type of machine learning approach that learns the underlying patterns or distributions of a dataset and uses this information to generate new, similar data points. It captures the probabilistic structure of the input data.

##### Variational autoencoder (VAE)

A variational autoencoder is a machine learning framework designed to encode high-dimensional data into a compact latent space while ensuring the ability to reconstruct the original data. It combines probabilistic modelling with neural networks for both compression and data generation.

##### Spurious correlation

A spurious correlation refers to an apparent association between two variables that arises due to a hidden factor or random chance, rather than a true causal relationship. This can lead to misleading patterns in machine learning models.

##### Invariant representation

An invariant representation isolates essential features of data that remain consistent across different conditions or transformations. It helps in focusing on meaningful signals by ignoring irrelevant variations, such as noise or confounding factors.

##### Latent variables

Latent variables are hidden or unobserved variables that are not directly measurable but influence the observed data. In machine learning, they represent underlying factors or structures inferred by the model to explain patterns in the data. For example, in generative models like variational autoencoders, latent variables help encode compressed representations of the data, capturing essential features or variations.

##### Prior distribution

A prior distribution in machine learning represents initial beliefs or assumptions about the distribution of a variable before observing any data. In generative models, like variational autoencoders, the prior is often a distribution like Gaussian used to regularise the latent variables.

##### Training data

Training data is the labelled or unlabeled dataset used to fit and optimise a machine learning model. The model learns patterns, relationships, and representations from this data to perform a specific task.

##### Testing data

Testing data is an independent dataset used to assess the performance of a trained machine learning model. It provides a measure of how well the model generalises to unseen data.

##### Generalizability

Generalizability in machine learning refers to the ability of a trained model to perform well on new, unseen data (testing data) that was not part of its training set. It indicates how effectively the model has learned patterns that are broadly applicable rather than memorising specific details of the training data. Generalizability is crucial for ensuring the model’s reliability in real-world applications.

The inputs (*X*, *B*, *T*) are encoded into two distinct latent variables, *Z_I_* and *Z_S_*, via a nonlinear encoder (Figure 1a). The latent variables *Z_I_* encode an invariant representation of cells conditioned on the biological variables *B*, while *Z_S_* models a spurious representation that captures unwanted variability in the data (Figure 1b). *Z_I_* is explicitly designed to be invariant to technical artefacts and is disentangled from *Z_S_* (details provided in the Methods section). In the context of large datasets with diverse biological and technical covariates, *Z_I_* represents a batch-corrected embedding of the data that identifies cell states faithfully and is suitable for downstream analyses, including clustering, cell annotation, and differential abundance testing (Figure 1b).

The learned prior distributions can be used to infer relationships across biological conditions or samples. The trained model can be applied to map new datasets onto a reference atlas, with built-in classifiers available for cell annotation in these new datasets. **c,** inVAE enables several applications, including the identification of cell states across different health and disease conditions, developmental stages, or spatial cell locations.

The latent variable *Z_S_* retains information related to spurious correlations typically regressed out during batch correction. We assume that the spurious correlations stemming from data biases are unrelated to the causal explanation of interest, although they are required for a perfect reconstruction of the input data by the model. By learning *Z_S_*, the inVAE model enhances interpretability, allowing visualisation of the data biases (Figure 1b). Analysing *Z_S_* provides insights into the influence of technical artefacts on sample proximity. Moreover, the independence of *Z_S_* from *Z_I_* serves as an indicator of model performance, guiding potential adjustments to hyperparameters. To reconstruct the input, a non-linear decoder maps the latent variables *Z* = (*Z_I_*, *Z_S_*) into the parameters of a negative binomial distribution, which is particularly well-suited for modelling count data frequently encountered in genomic applications (Figure 1a).

In order to model the distributional shifts caused by biological or technical factors, inVAE infers conditional prior distributions over both invariant and spurious latent variables (Figure 1a, right). To learn these priors, inVAE utilises Gaussian distributions, directly estimating the means and variances of the distributions as a function of biological (*B*) or technical covariates (*T*) implemented by two separate neural networks. The parameters of these networks are jointly optimized with the rest of the model using backpropagation and the designated loss function. The means of the prior distributions can be further used to derive, for example, the severity across disease conditions or clustering datasets based on their technical artefacts (Figure 1b). Using inVAE, cells from diverse conditions are integrated into a shared phenotypic latent space, and by aligning the latent variable distributions with the conditional prior distributions through Kullback-Leibler (KL) divergence, we capture both common and condition-specific cell states within the invariant representation (details provided in the Methods section).

To process a new dataset, reference mapping and label transfer are performed by freezing the weights of the encoder and first inferring the latent representation by reusing previous knowledge about known covariates. For label transfer, a separate classifier is then trained on the invariant latent representation of the new dataset to predict unlabelled cells (details are provided in the Methods section).

In summary, inVAE provides a shared representation of multivariate datasets, incorporating prior knowledge such as metadata, into the integration process to accurately identify cell states. inVAE improves interpretability by dissecting the biological and spurious sources of variation.

### inVAE accurately identifies disease-specific cell states in human heart failure

Recent single-nucleus RNA sequencing (scRNA-seq) analyses of healthy and diseased adult human hearts provide an opportunity to explore disease-specific molecular changes at the cellular level^5,32–34^. Although pathogenic variants have been identified in genetic cardiomyopathies like dilated cardiomyopathy (DCM) and arrhythmogenic cardiomyopathy (ACM)^35^, the shared and unique molecular signatures of the distinct genetic cardiomyopathies in heart failure remain poorly understood. To facilitate such discoveries, comparing the impact of different pathogenic variants is essential^32^. However, integrating and identifying cellular populations across samples from individuals of different age, sex, and unknown disease or genetic background poses significant challenges.

To assess the capacity of inVAE to link molecular to clinical traits, we used a large dataset of human heart failure^32^. We specifically reanalysed snRNA-seq data from explanted ventricular tissues of patients with pathogenic variants in genes related to DCM (*TTN*, *LMNA*, and *RBM20*) and ACM (*PKP2*) (n=16) as well as controls (n=6). We utilised pathogenic variants and anatomical regions (left and right ventricle) as biological covariates (see Methods). Our final dataset included 284,727 high-quality nuclei from both diseased and healthy left and right ventricles from male and female donors aged 30-60 (Figure 2a).

**Figure 2:**
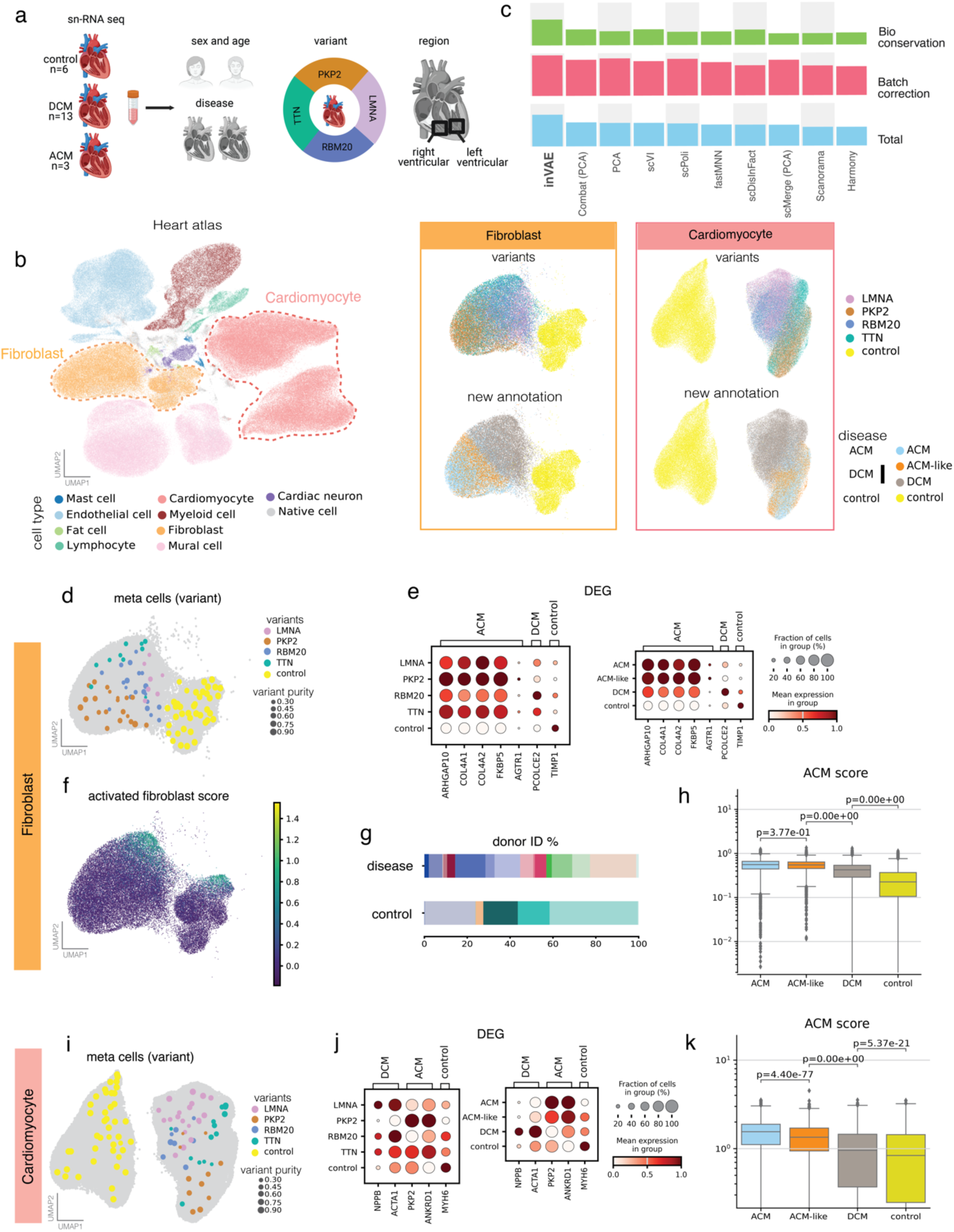
inVAE reveals disease-specific cell states in human heart failure. **a**, The dataset encompasses samples from control and patients with dilated cardiomyopathy (DCM) and arrhythmogenic cardiomyopathy (ACM), with diverse pathogenic variants, as well as donors of varying sex and age. **b,** The Uniform Manifold Approximation and Projection (UMAP) visualisation of inVAE’s invariant representation highlighted by cell types and disease-specific states, focusing particularly on cardiomyocytes and fibroblasts. **c,** Overall biological conservation and batch correction scores across leading integration methods using scIB metrics^41^. **d,** Meta-cell analysis using seaCell reveals variant-specific regions in the fibroblast cell embedding. Each dot represents a meta cell, which is a cluster of neighbouring cells, and is coloured according to the most frequent variant observed within that meta cell. The size of the dots indicates the percentage of cells within the meta cell that are labelled with the top variant. Meta cells are projected on the UMAP representation of fibroblast cells, shown in grey. **e,** Dot plot displaying log2-transformed expression of differentially expressed genes (DEG) across variants and disease conditions in fibroblasts. **f,** Activated fibroblast scores using *POSTN* and *FAP* genes. **g,** Sample compositions for cells with high fibroblast score (>1.0) for both diseased (ACM and DCM) and control cells. **h,** The box plot showing the distribution of ACM scores using *ARHGAP10*, *COL4A1*, *COL4A2*, and *FKBP5* genes across disease states for fibroblasts. **i,** Meta cell analysis identifying variant-specific regions in the embedding of cardiomyocytes. **j,** Dot plot displaying DEG across variants and disease conditions in cardiomyocytes. **k,** Box plot showing the distribution of ACM score using PKP2 and ANKRD1 genes across disease states for cardiomyocytes.

The manifold generated by inVAE accurately identified all cell types previously detected in the original article (Figure 2b), and displayed a clear separation between control and disease samples (Supplementary Figure 1). inVAE outperformed 9 alternative data integration methods (scVI^19^, Combat-PCA^36^, PCA, scPoli^26^, fastMNN^21^, scDisInFact^37^, scMerge-PCA^38^, scanorama^18^, harmony^22^) in preserving biological variation between diseased and control samples while maintaining comparable performance in batch correction (Figure 2c, see Methods). Conventional metrics for evaluating batch correction usually focus on merging samples from multiple batches. This poses a problem in datasets with multiple diseases, as disease-specific cell states should remain distinct (and not integrated). Thus, what is considered by such methods as “successful batch correction” could risk over-correcting biological differences. To accurately assess the robustness of the inVAE representation of the heart disease dataset, we conducted a detailed analysis of the inferred cell states.

The visualisation of our embedding revealed distinct distributions of fibroblasts and cardiomyocytes that were associated with specific genotypes (Figure 2b), unlike in the original study^32^, where the manifold masked these associations. We examined if the distribution of cells in inVAE’s manifold revealed known molecular signatures associated with the disease. For this, we first generated meta cells via seaCell^39^ using inVAE’s embedding of fibroblasts, and identified neighbourhoods enriched for specific genotypes and control samples (Figure 2d). ACM cells (*PKP2* variants) were located in a specific region of the embedding, while DCM variants (*LMNA*, *TTN*, and *RBM20*) showed higher similarity to each other, resulting in more merged samples. This was further supported by the expression of known genotype-specific genes from the literature, such as the upregulation of *AGTR1* in PKP2 cells, the variable expression of *ARHGAP10*, *COL4A1*, *COL4A2*, and *FKBP5* in all disease samples, and the downregulation of *TIMP1* in all disease samples (Figure 2e). Thus, this new embedding successfully highlighted disease-specific differences amongst cells.

We next investigated whether inVAE could reveal new cell states. We identified a distinct fibroblast cell state present in both disease and control samples, expressing activated fibroblast markers *POSTN* and *FAP*^33,40^ (Figure 2f), which was not identified in the original study. These activated fibroblasts were derived from all donors (Figure 2g), suggesting that inVAE successfully corrected for batch effects by aggregating activated fibroblasts from control and disease samples.

Additionally, within the representations of both fibroblasts and cardiomyocytes, we identified subsets of cells from DCM hearts (termed DCM cells) that were co-located in the embedding with ACM cells (Figure 2b, we termed these cells ACM-like fibroblasts and ACM-like cardiomyocytes). This suggests that cellular alterations in DCM can resemble ACM changes.

To further understand the molecular identity of these newly identified cell types, we first focused on investigating the new fibroblast subsets associated with disease genotype. The ACM-like fibroblasts had an enrichment score for ACM markers (*ARHGAP10, COL4A1, COL4A2*, and *FKBP5*), and a significant overlap in the transcriptomic signatures with ACM cells (Figures 2e and 2h, Supplementary Figure 2a). ACM-like fibroblast cells were present across all DCM samples, with most of these samples derived from *TTN* and *RBM20* variants (Supplementary Figure 2b). Pathway enrichment analysis of genes differentially expressed in DCM compared to ACM-like fibroblasts revealed significant enrichment in pathways related to extracellular matrix and collagen organisation, consistent with known transcriptional changes in DCM fibroblasts associated with extracellular matrix and fibrotic progression^42^ (Supplementary Figure 2c-d).

Secondly, we focused on the new cardiomyocyte subsets associated with disease genotype. Using meta cells within the cardiomyocyte embeddings, we identified neighbourhoods significantly enriched for control, *LMNA* (a causal gene of DCM), and *PKP2* (a causal gene of ACM) expressing cells (Figure 2i). Consistent with previous reports^32^, DCM upregulated *NPPB* and *ACTA1*, ACM upregulated *PKP2* and *ANKRD1*, while control cells upregulated *MYH6* (Figure 2j). We then calculated enrichment scores for ACM markers (*PKP2* and *ANKRD1*), revealing distribution shifts across ACM, DCM, and control cardiomyocytes, and highlighting the similarity between ACM and ACM-like cardiomyocytes, consistent with our earlier findings (Figures 2j and 2k). Differential expression analysis showed a strong overlap of differentially expressed genes between ACM and ACM-like cardiomyocytes (Supplementary Figure 3a), further demonstrating the robustness of inVAE’s representation. Sample composition indicated that the ACM-like population includes cardiomyocyte cells from all DCM samples, with a predominance of cells harbouring the *TTN* variant, known for being less severe and more responsive to standard therapy than other forms of DCM^43^ (Supplementary Figure 3b). Pathway enrichment analysis of differentially expressed genes in ACM-like cardiomyocytes showed significant enrichment in mitochondrial function and oxidative phosphorylation, indicating relatively preserved metabolic function in ACM-like cells compared to DCM cells (Supplementary Figure 3c-d). Collectively, these findings suggest that ACM-like cardiomyocytes share phenotypic traits with ACM cardiomyocytes and are marked by preserved metabolic function and lower stress-related gene expression compared to other DCM cells.

In summary, inVAE characterises disease-specific cell states including the phenotypic features of distinct genotypes, while capturing known markers and pathways associated with poor outcomes.

### inVAE offers enhanced interpretability by accurately disentangling biological signals from noise and generalises to unseen data of human heart failure

To demonstrate inVAE’s interpretability and generalizability in a comparable context, we generated a comprehensive atlas of human heart failure and used it to predict disease and cell types in unseen samples, allowing for direct benchmarking against specialized integration methods. We collected and harmonised the annotations of three previously published datasets from 110 donors across three conditions: control, DCM and hypertrophic cardiomyopathy (HCM)^32–34^ (see Methods). To evaluate inVAE’s generalizability as a reference model for label transfer, we first split the data into training and testing data. We then built an integrated atlas of the heart using the training data and transferred the labels to the testing data after projecting it on the reference atlas.

The training and testing datasets were sourced from different studies, ensuring no overlap in donor IDs (Figure 3a). Two studies containing both control and DCM, from Reichart et al.^32^ and Kanemaru et al.^33^, were used for training. A third study, from Chaffin et al.^34^ originally containing control, DCM and HCM samples, served as the testing set. We further split the testing data into out-of-sample and out-of-disease to study two different scenarios. In the out-of-sample testing data, the disease conditions matched those in the training subset (i.e. we only considered control and DCM data from Chaffin et al.) (scenario 1, Figure 3a). In contrast, the out-of-disease testing data consisted only of HCM donors from Chaffin et al. (scenario 2, Figure 3a). We anticipated that both scenarios would be challenging due to batch effects expected from different studies, with the out-of-disease scenario (scenario 2) being particularly difficult due to potential biological variation introduced by differing disease conditions.

**Figure 3:**
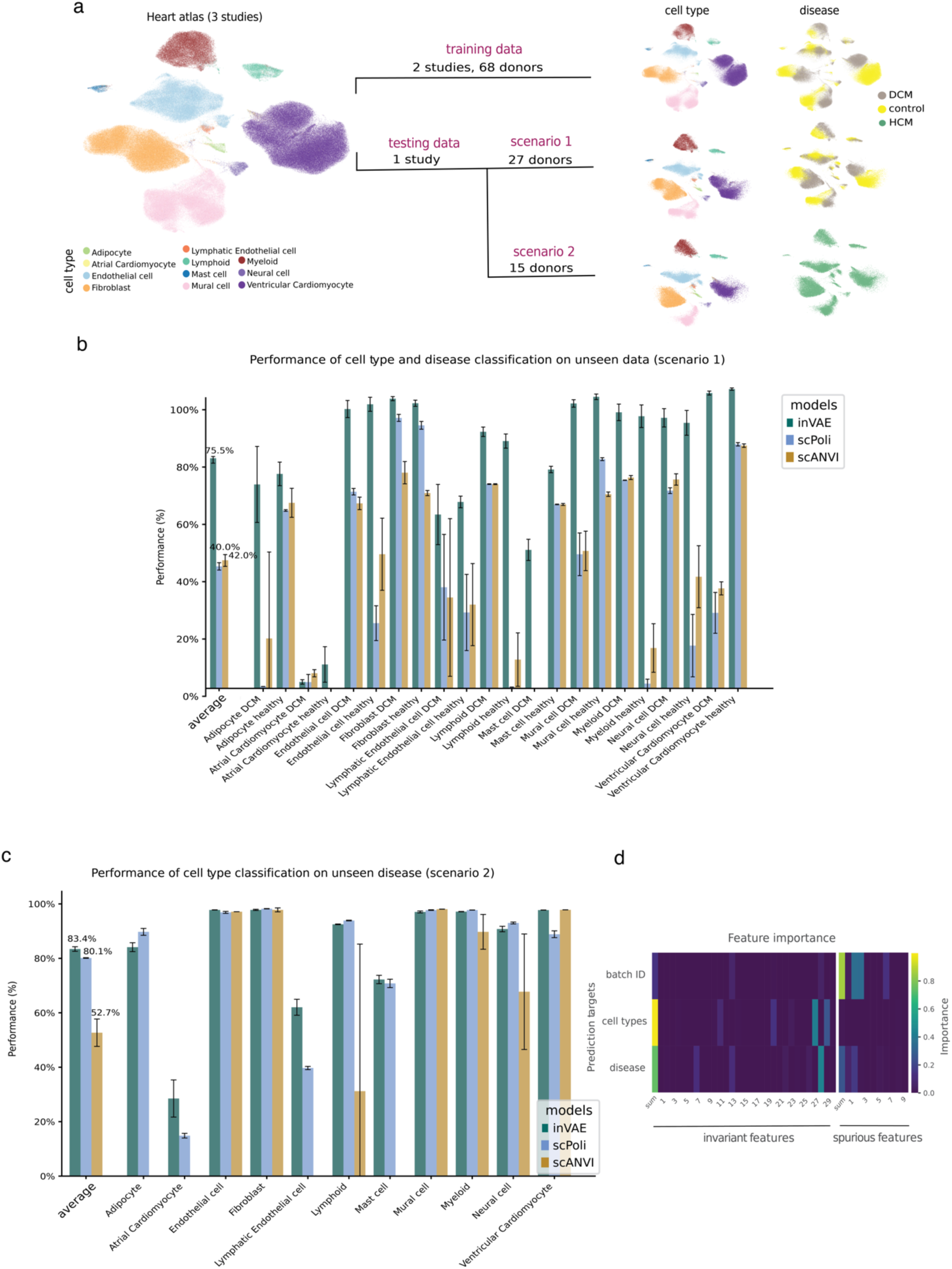
inVAE transfers labels and disease states in the human heart to an unseen study. **a**, The UMAP visualisation of inVAE’s extended heart atlas. The inVAE model is trained on two studies comprising samples from control and DCM donors. This trained model is then applied to map an unseen study onto the reference atlas. Two scenarios are considered: in the first, the unseen study involves similar conditions (control and DCM) to the training data, while in the second, it includes an unseen disease (HCM). **b,** Bar plot indicating the performance of disease and cell type classification for scenario 1. Shown are the average (Macro F1 scores) and per cell type and disease prediction performance (F1 scores). The prediction uncertainty is displayed per class. **c,** Cell type classification performance for scenario 2. Overall prediction performance (Macro F1 scores) and cell-type specific performance are shown (F1 scores). **d,** The importance of latent variables for different classification tasks (batch ID, cell type and disease) using a permutation-based importance measure metric^44^. ACM, arrhythmogenic cardiomyocyte; DCM, dilated cardiomyocyte; HCM, hypertrophic cardiomyopathy.

To assess inVAE generalizability for label transfer, we first trained the model exclusively on the training data (from Reichart et al.^32^ and Kanemaru et al.^33^), creating a joint embedding of control and DCM cells. The integration results demonstrated a clear separation between control and disease cells (Figure 3a, right top row). Next, we trained a classifier using the invariant latent representation from the training data and tested the two scenarios. In the first scenario (out-of-sample), where the testing data shares similar conditions with the training data, we used the classifier to predict both cell type and disease status for each cell. The results showed a significant improvement with inVAE (Figure 3b), enhancing its generalizability for label transfer to unseen data. In the second scenario, which involved samples from an unseen disease (out-of-disease), we trained the classifier to predict only cell type labels using the learned latent representation. Cell type prediction is generally easier than disease prediction due to the presence of strong signals distinguishing cell types. The results indicated that inVAE consistently improved cell type classification accuracy, outperforming related methods (Figure 3c).

We expected the invariant and spurious latent spaces to be effectively disentangled with minimum shared information. In inVAE’s framework, the invariant representation should capture biological signals, while the spurious representation encodes technical variations. To assess this separation, we trained different classifiers using both invariant and spurious latent variables to predict targets such as donor ID, disease condition, and cell type. A permutation-based importance metric^44^ assessed the contribution of each latent variable to each prediction target (Figure 3d, Methods). In this analysis, high variable importance suggests relevance to the target prediction. The results revealed that cell type and disease predictions relied primarily on invariant variables, with spurious latent variables having near-zero importance. As expected, donor ID prediction depended mainly on spurious variables. These findings demonstrate that inVAE effectively separates batch effects from biological variation. They also highlight inVAE’s interpretability, which allows validation of this disentanglement—a feature often missing in other batch correction methods that do not explicitly model or remove spurious variations.

In summary, inVAE significantly outperforms existing methods in disease and cell type classification, effectively creating representative atlases that capture true biological variation while filtering out technical artefacts. Moreover, inVAE offers a unique advantage by enabling the analysis of the spurious latent space, enhancing model interpretability by pinpointing technical sources. For downstream analyses that rely on invariant representations, the spurious representations provide valuable insights into the excluded information and its dependency on covariates, enriching the overall understanding of the data.

### inVAE identifies temporal cell states in human fetal lung

Accurate modelling of differentiation trajectories in dynamic biological systems requires time-course single-cell analyses, which capture snapshots of consecutive developmental states. However, integrating these datasets poses challenges due to technical and temporal variations, and existing methods often struggle to account for cellular dynamics. We next used inVAE to infer a representation of real dynamic systems that maintains the developmental stage of cells.

We applied inVAE to a temporal single-cell transcriptomics atlas of the human fetal lung^45^, which includes 65,564 cells from 29 donors and 8 time points spanning 5–22 post-conception weeks (PCWs) (Figure 4a, Methods). The integrated single-cell atlas using inVAE successfully identified all 144 cell states previously characterised in the original study using principal component analysis^45^ (Figure 4b, Supplementary Figure 4). We also integrated the data using 4 additional methods (scVI, Combat-PCA, PCA, scPoli) and observed an improvement in batch effect correction and bio-conservation (Figure 4c).

**Figure 4:**
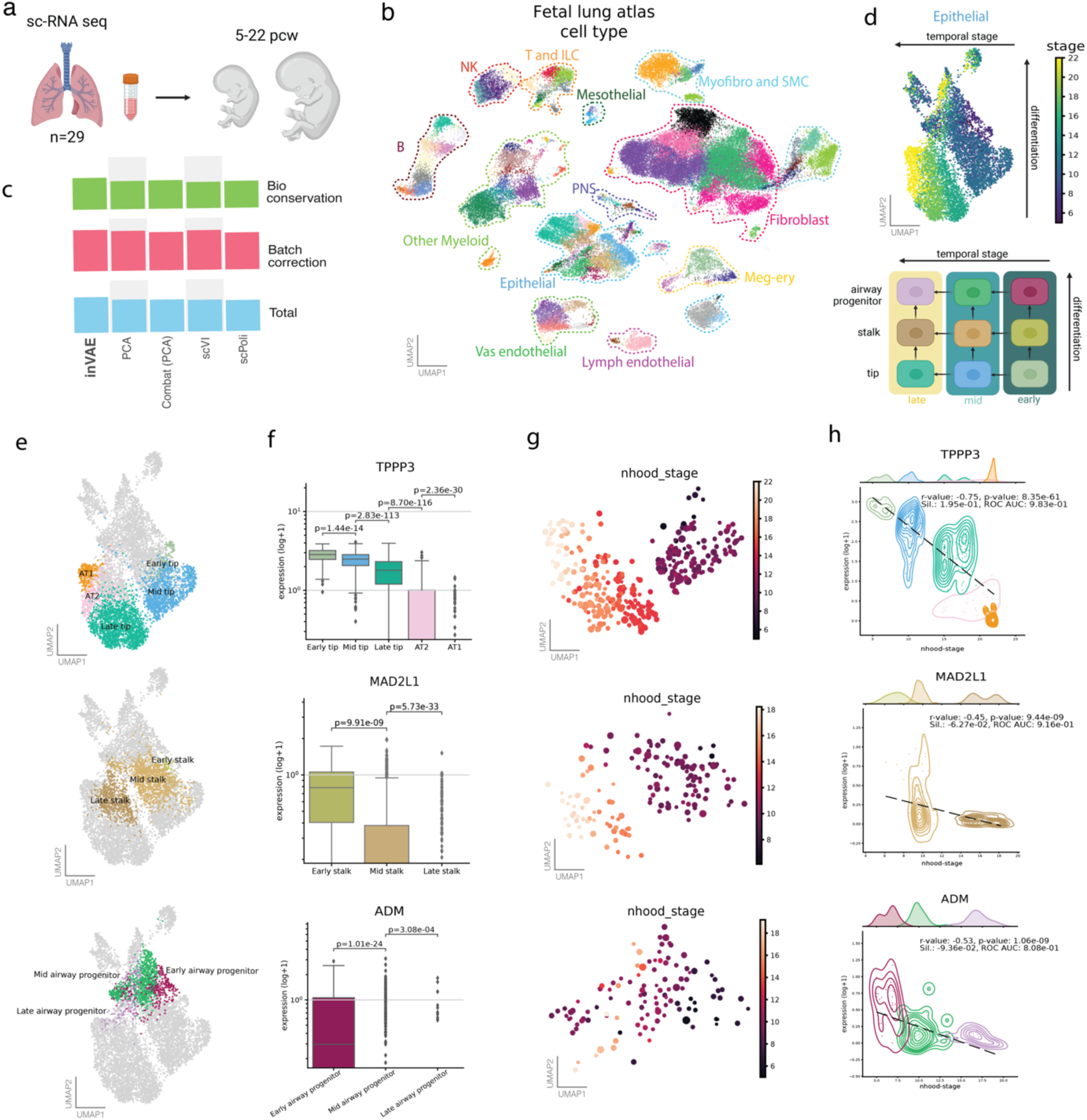
inVAE identifies temporal cell states in the human fetal lung atlas. **a**, The experimental design of the fetal lung atlas study using samples from 29 donors, spanning 5 to 22 post-conception weeks (PCW). **b,** The UMAP visualisation of the atlas after applying the inVAE integration method coloured by cell type. Major cell types are labelled. **c,** Overall biology conservation and batch correction scores across the top five integration methods using scIB metrics^41^. **d,** top: UMAP focused on epithelial cells coloured by developmental stage. Shown are differentiation and temporal state axes. Bottom: The schematic overview of the cell types and their transitions along the two axes. **e,** UMAPs revealing similar cell types at three developmental stages, assigned as early, mid and late. **f,** The expression patterns of key markers (*TPPP3*, *MAD2L1*, and *ADM*) at each developmental stage shown using box plots across cell types. **g,** Neighbouring cell relationships depicted using Milo^46^. Each dot in the plot represents a neighbourhood of cells, coloured by the average developmental stage of the cells within that neighbourhood (termed ‘nhood-stage’). **h,** The distribution of marker genes in each cell type as a function of nhood-stage. The dashed lines show the correlation between the gene expression and nhood-stage by fitting a linear model. NK, natural killer cell; B, B cell; T, T cell; ILC, innate lymphoid cell; Myofibro, myofibroblast cell; SMC, smooth muscle cell; Meg-ery, megakaryocyte and erythroblast cells; PNS, peripheral nervous system; sil., silhouette; ROC AUC, Receiver Operating Characteristic Area Under the Curve.

In the fetal lung, epithelial cells progress through differentiation from tips to stalks to airway progenitors, with additional temporal variations influencing each stage (Figure 4d, bottom). This means that at each temporal stage–early (5–6 PCW), mid (9–11 PCW), and late (15–22 PCW)–the identity of tips, stalks, and airway progenitors evolve. By including developmental stages as biological covariates, inVAE accurately captured these temporal variations among epithelial cells while preserving their developmental trajectories (Figure 4d-e). In the inVAE manifold, cell states follow a developmental trajectory along the vertical axis (from tip to stalk to early airway) and reveal temporal state changes along the horizontal axis (from early to mid to late) (Figure 4d, top). These dynamic cell transitions along the horizontal axis were not captured in the original manifold, where cells were grouped as distinct populations by their stage (early, mid, and late).

Stage-specific markers (*TPPP3, MAD2L2, and ADM*) showed a gradient of change across the early-mid-late axis (Figure 4f). To quantify the accuracy of the cellular transition within the latent representation, we applied Milo^46^, which assigns cells to partially overlapping neighbourhoods on a k-nearest neighbour graph (Figure 4g). This approach, based on cell-cell similarity within inVAE’s latent space, allowed us to assess the model’s representation. Results showed a strong correlation between the mean expression of key markers and neighbourhood-stage (Figure 4h), confirming inVAE’s ability to capture both temporal and lineage-specific transitions in cell states.

Additionally, we identified two distinct cell states with unique transcriptomic signatures in late tip cells at 15 and 18 PCW, which were not observed in the original study (Supplementary Figure 5a). Using scFates^47^ on inVAE’s latent embedding, we inferred the pseudo-ordering of cells along the tip axis and identified genes with dynamic changes throughout this transition (Supplementary Figure 5b). The analysis revealed distinct gene modules associated with each stage of the trajectory, which were also differentially expressed between the two late tip cell states (Supplementary Figure 5b). Among these gene modules, we highlight the expression patterns of three genes with distinct expression patterns: *TPPP3*, *CD36*^48^, and *EEF1G* (Supplementary Figure 5c).

Airway progenitor and stalk cells differentiate into either pulmonary or *GHRL*+ neuroendocrine cells as described by He et al.^45^ (Supplementary Figure 6a). Using PAGA trajectory inference on inVAE’s latent embedding, we accurately derived these developmental trajectories, which were further validated by the expression of specific markers, including *ASCL1*, *MAD2L1*, *NEUROD1*, *NKX2-2*, and *NEUROG3*^45^ (Supplementary Figure 6b-d). Thus, the inVAE’s embedding captures final cell states and, in addition, suggests that the starting population previously defined “as airway progenitors and stalk cells” may already be developmentally skewed to distinct fates, with airway progenitors more closely linked to *GHRL*+ NE and stalk cells to pulmonary NE cells (Supplementary Figure 6c).

In conclusion, inVAE successfully identified high-resolution cell states in the fetal lung, derived both developmental trajectories and temporal cell states in epithelial cells, and suggested an earlier cell fate split in the neuroendocrine developmental trajectories.

### inVAE unravels spatial cell-state variability in adult human lung

We next applied inVAE’s to disentangle spatially resolved cell states using a high-resolution single-cell atlas of the healthy human lung^49^, which encompasses data from five anatomical locations along the proximal-to-distal axis (Figure 5a). This atlas is comprehensively annotated with 78 distinct cell types derived from single-cell and single-nucleus RNA sequencing of the trachea, two bronchi, and upper and lower parenchyma regions.

**Figure 5:**
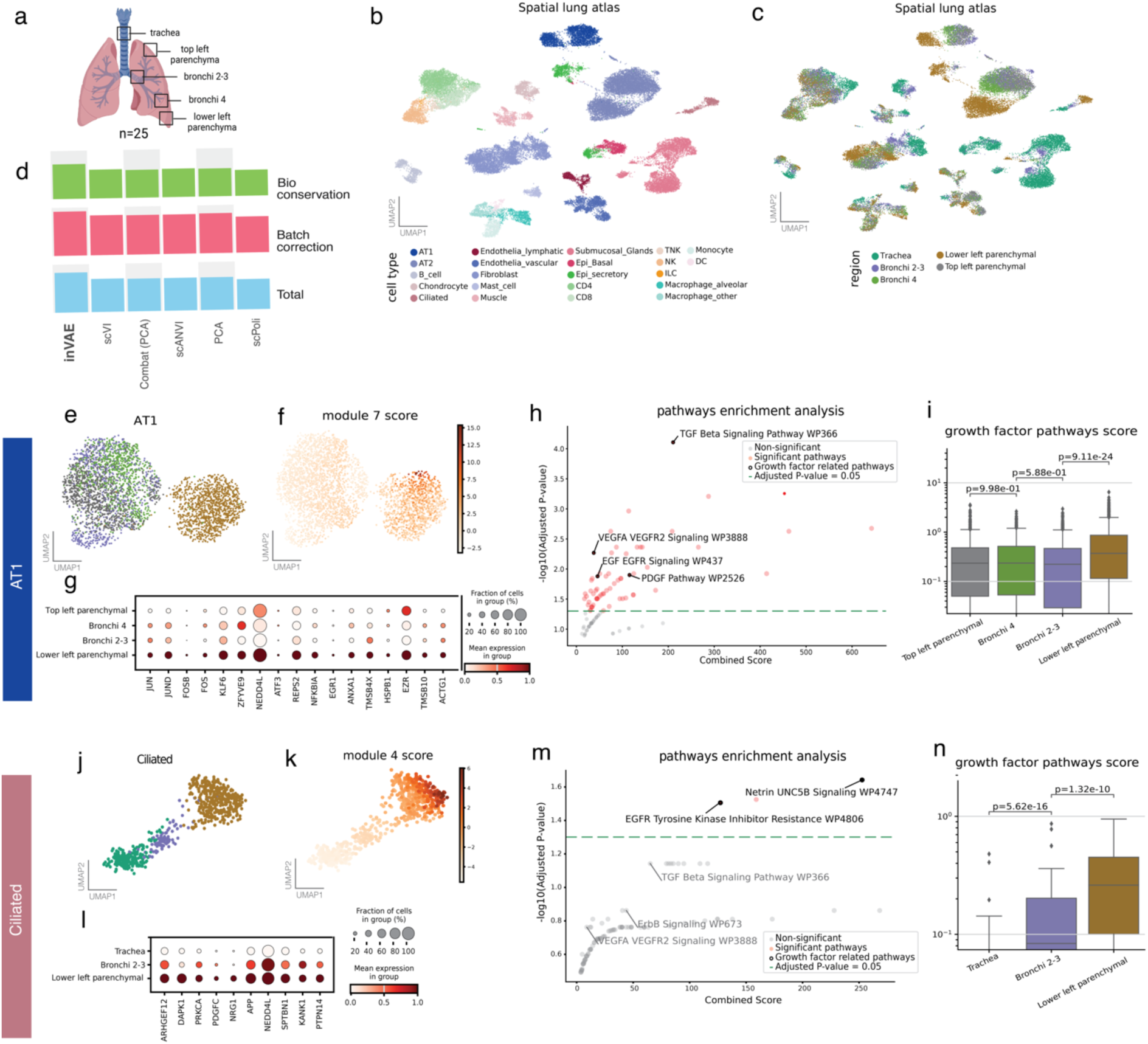
inVAE identifies spatial cell states in the adult human lung atlas. **a**. Lung samples collected from multiple anatomical regions of the lung. **b-c.** Uniform Manifold Approximation and Projection (UMAP) visualisations of the atlas after inVAE integration highlighted by cell types and regions. **d.** Overall biology conservation and batch correction scores comparing top-performing integration methods using scIB metrics^41^. **e.** A UMAP visualisation highlights AT1 cells, coloured by anatomical region. **f.** Enrichment score for differentially expressed genes (DEGs) in the lower left parenchyma, identified using unsupervised Hotspot. **g.** A dot plot displaying the log2-transformed expression of the DEGs. **h.** Enriched pathways, ranked by their –log10-transformed adjusted p-value. The dashed line marks significantly enriched pathways (p-value > 0.05). **i.** The enrichment score for growth factor pathways in each AT1 cell region. **j.** UMAP visualisation highlights ciliated cells, coloured by anatomical region. **k.** The enrichment score for DEGs of the lower left parenchyma, identified by unsupervised Hotspot. **l.** A dot plot showing the corresponding DEGs and their log2-transformed expression. **m.** Enriched pathways in the lower left parenchyma of ciliated cells. **n.** The enrichment score for growth factor pathways in each region of ciliated cells. AT1, alveolar type 1.

By training inVAE with anatomical regions as biological covariates, we captured high-resolution spatial variation in cell type distributions, achieving greater bio-conservation and better batch correction than conventional methods (Figures 5b-d). inVAE highlighted unique spatial and functional characteristics in alveolar type 1 (AT1) cells, responsible for gas exchange (Figures 5e), and in the ciliated cells lining the airways (Figures 5j). Using Hotspot^50^ we identified key pathways and gene modules enriched in specific spatial regions (see Methods).

In AT1 cells, we identified eight gene modules with distinct spatial patterns (Supplementary Figure 7a). The lower parenchyma was particularly distinct from other lung regions, with module 7 showing the strongest association with this area (Figure 5f). In the experimental design of this study each sequencing pool contained samples from each region, but with each region being derived from a different donor. Supplementary Figure 8a shows that the high contribution of module 7 to the lower lobe signature was consistent across all donors analysed, and did not track with the sequencing pool. We used gene autocorrelation metrics—Z-scores and p-values—to find the expression patterns among the top differentially expressed genes (Figure 5g). Pathway enrichment analysis identified significant pathways, including TGF-beta^51^, EGF-EGFR^52^, PDGF^53^, and VEGFA-VEGFR2^54^, known to contribute to lung fibrosis, a condition that often begins in the lower lobes^55^ (Figures 5h–i).

Ciliated cells exhibit spatial differences across lung regions, leading to functional variations^56,57^. Using inVAE’s embedding, Hotspot identified five gene modules for ciliated cells (Supplementary Figure 7b), with module 4 highly expressed in the lower parenchyma (Figure 5k), and unbiasedly distributed across donors and sequencing pools (Supplementary Figure 8b). Enrichment analysis revealed that ciliated cells in the lower left parenchyma are primarily enriched in netrin signalling and other growth factor related pathways (Figures 5m-n)^58^. Netrin-1 has been linked to pulmonary fibrosis and interstitial lung disease rheumatoid arthritis patients^59^. While previous studies have provided limited insights into the regional variation of ciliated cells, our analysis identifies specific gene modules and signalling pathways associated with distal lower bronchi, offering new perspectives on their function and regulation throughout the lung.

In summary, inVAE uncovered not only spatially-resolved cell states but also preserved key gene expression patterns unique to specific lung regions, demonstrating its ability to highlight spatially significant variations across cell types.

## Discussion

inVAE is a generative representation learning model designed to incorporate biological covariates, infer conditional cell distributions, and retain essential and subtle biological variations. inVAE represents the latent space with two sets of variables: invariant and spurious. By disentangling these two representations, it enhances the interpretability of the latent space by dissecting the sources of variation in the data. The invariant representations are suited for downstream analyses like high-resolution cell state identification, while spurious representations capture the distribution of biases across cells or samples. inVAE infers prior distributions of cells specific to each biological condition, like disease variants, or technical biases like study. This provides insights into both biological and technical variation across samples with potential implications for understanding pathogenesis, identifying biomarkers, and discovering drug targets^10,12,33,60–62^. While invariant representation models^63–66^ are known to generalise well to unseen datasets generated under similar conditions, inVAE transfers labels to new datasets and, crucially, identifies novel cell states when the cells in a new dataset undergo domain shifts, such as those caused by previously unseen diseases.

We validated inVAE’s capabilities through four complex data integration tasks aimed at identifying cell states from samples with diverse biological conditions. In the first task, we generated a human heart atlas encompassing both healthy and diseased samples, showing that inVAE could stratify donors based on the genetic impact of pathogenic variants with higher resolution than previous methods. For the second task, we expanded this atlas to include samples from three distinct studies, demonstrating the model’s ability to generalise to unseen data, including samples from previously unobserved disease conditions, with significant improvements in cell and disease annotations over leading methods. In the third task, inVAE successfully captured temporal cell states and inferred realistic temporal cell state relationships from human lung developmental data. Future experimental work will be required to validate these suggested cell states in vivo. In the final task, inVAE identified spatial cell states in spatially-resolved datasets of human lung cells, and highlighted gene modules and signalling pathways that reflect regional variations of AT1 and ciliated cells. Identified gene modules were linked to fibrosis, recapitulating the known spatial distribution of this disease. Collectively, inVAE allowed for the identification of new cell states, more precise pathway enrichment, and enhanced differential expression gene analysis through improved sample stratification.

In this manuscript, we demonstrated the success of inVAE in integrating large cohorts using categorical covariates such as disease variants modeled with one-hot encoders. However, some real-world applications might involve continuous covariates, such as spatial positions of cells along a spatial axis. Although inVAE supports continuous covariates through multi-layer perceptron (MLP) encoders, we have not extensively tested this capability, leaving it as an interesting direction for future work. Additionally, the one-hot encoding of categorical covariates treats them as independent variables, with equal distances between encoded categories. While the model effectively infers the similarities and differences across prior distributions, it could benefit from incorporating hierarchical structures or prior knowledge about the relationships between covariates. For instance, in disease modeling, where certain variants may share closer biological similarities, leveraging this information could reduce the search space and accelerate convergence. However, integrating such prior knowledge would require reliable biological metrics to assess these relationships, offering an exciting avenue for future exploration.

While our current evaluation focuses on single-cell transcriptomics, we envision future applications of inVAE in integrating other modalities. In addition to the negative binomial distribution, Poisson and Gaussian distributions are already implemented, enabling the integration of scATAC-seq and spatial data. Another promising future direction for inVAE is the identification of key genes—whether causal, condition-specific targets, or those predominantly influenced by noise—providing deeper insights into disease mechanisms.

In conclusion, inVAE enables comprehensive analysis of multivariate data while disentangling true biological signals from technical biases. We anticipate that inVAE will play a key role in large-scale atlasing efforts, helping to stratify patients based on various biological factors and across a range of diseases.

## Methods

### Ethics statement

This study relies on the analysis of previously published data, which were collected with written informed consent obtained from all participants and comply with the ethical guidelines for human samples.

### Data preprocessing

The inVAE models were trained using raw counts. Further details are explained in the following.

### Reichart et al. heart data preprocessing

The raw gene expression data was downloaded from Reichart et al.^32^. For this study, we selected samples from control, TNN, LMNA, RBM20, and PKP2 donors aged 30–60. This preprocessing resulted in a final dataset of 284,727 cells and 5,000 highly variable genes, which were used for training.

### Heart atlas data curation

We collected the raw gene expression counts from three studies—Kanemaru et al. (2023), Reichart et al.^32^, and Chaffin et al.^34^. We randomly selected 400,000 cells due to resource constraints. We used Scanpy package^67^ to filter out low-quality cells, removing those with counts lower than 500 and higher than 15,000. Doublets were removed using Scrublet^68^, retaining cells with a doublet score below 0.2. We trained the models using 4,000 highly variable genes and 167,633 cells in total. The annotations were harmonised across the three studies using celltypist^69^.

### Splitting heart atlas data into train, test, and validation

The filtered heart atlas data was divided into training and validation sets by randomly selecting 90% (n=167,633) and 10% (n=18,625) of the cells from both the Kanemaru et al.^33^ and Reichart et al.^32^ studies, respectively. For scenario 1, the test set consisted of cells (n=84,801) from the Chaffin et al.^34^ study under the same conditions as the training and validation sets, specifically control and DCM. In scenario 2, the test set included cells (n=52,270) from the Chaffin et al. study collected from donors with HCM, representing an unseen condition during training.

### Fetal lung data preprocessing

We re-analysed the fetal lung atlas in He et al.^45^ using the set of highly variable genes (n=5101) identified in the original study. To refine this gene set, we removed a cluster of genes (n=121) associated with cell cycle activity (e.g., CDK1, MKI67, CCNB2) by applying Leiden clustering^70^ in PCA space (n_pcs=15) of the genes using Scanpy. This preprocessing resulted in a dataset comprising 65,564 cells and 4,980 highly variable genes, which was subsequently used for training.

### Spatial lung data preprocessing

The raw gene expression data was downloaded from Madissoon et al.^49^. A total of 193108 cells and 4000 highly variable genes identified by the original paper were used for training.

### inVAE method

inVAE is a semi-supervised deep generative model that builds upon the framework of identifiable Variational Auto-Encoders (iVAE)^71^. The primary purpose of inVAE is to disentangle latent factors into two components: an invariant latent representation and a spurious one. This allows the model to distinguish signals originating from biological variations from those unwanted, like technical noise.

### Notations

In inVAE’s model, the variable 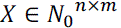 represents the observed data, such as gene expression counts in single-cell genomics, with *n* cells and *m* genes. The variable *B* ∈ *R ^n^*^×*b*^refers to biological or condition-specific covariates such as cell types or disease states, with *b* the number of dimensions resulting from concatenating one-hot encoded or continuous covariates. *T* ∈ *R^n^*^×*t*^ represents batch or technical covariates that may introduce unwanted variability, for example, sequencing technologies, with again *t* the number of dimensions resulting from concatenation. The latent representation is denoted by *Z* ∈ *R^n^*^×(*k*+1)^, which is further composed of two components: an invariant latent representation *Z_I_* ∈ *R*, and a spurious latent representation *Z_S_* ∈ *R^l^*., such that *Z* = (*Z_I_*, *Z_S_*). The component *Z_I_* represents the invariant latent representation, which encodes biological signals that are consistent across conditions, whereas *Z_S_* represents the spurious latent representation, which encodes technical noise or other unwanted variations. The approximate posterior distribution over the latent variables *Z*, given the observed variables *X*, *B*, *T*, is denoted by *q*_ϕ_(*Z*|*X*, *B*, *T*). This distribution is parameterized by a neural network with parameters ϕ. The prior distribution over the latent variables is denoted by *p_θ_*(*Z*|*B*, *T*), which can be further decomposed into the conditional priors *p_θ_*(*Z_I_*|*B*) and *p_θS_* (*Z_S_*|*T*). The likelihood of the observed data *X* given the latent variables *Z* is represented by **p*_f_*(*X*|*Z*), which is parameterized by a neural network with parameters *f*.

### Framework

inVAE encodes observed data *X*, along with biological covariates *B* and technical covariates *T*, into a structured latent space *Z* that consists of an invariant latent representation *Z_I_* and a spurious latent representation *Z_S_*. To achieve this, inVAE employs an encoder parameterized by a neural network that maps the inputs (*X*, *B*, *T*) to the approximate posterior distribution *q_ϕ_*(*Z* ∣ *X*, *B*, *T*). This distribution is modelled as a Gaussian distribution with a mean and variance parameterized by a neural network with parameters ϕ.

The latent representation *Z* is then used by the decoder to reconstruct the observed data. Specifically, the latent variables *Z* = (*Z_I_*, *Z_S_*) are passed through a decoder, which generates parameters for the data likelihood *p_f_*(*X*|*Z*). The data likelihood follows a Negative Binomial distribution, which is suitable for modelling count data and a popular choice for gene expression levels in single-cell genomics. In theory, any other sensible distribution could be used.

In the inVAE framework, two distinct conditional priors are assumed for the latent space. The first prior distribution is for the invariant latent representation and is denoted as *p_θI_*(*Z_I_*|*B*). This prior belongs to a general exponential family distribution and aims to capture dependencies that arise from biological mechanisms that remain invariant across different biological conditions. The use of an exponential family prior provides flexibility and allows the model to effectively represent complex interdependencies between invariant latent variables. The second prior distribution is for the spurious latent representation, denoted as *p_θS_*(*Z_S_*|*T*), and captures the technical variations that are specific to different conditions, such as batch effects or other forms of technical noise. In practice, both priors are modelled as Gaussian distributions with diagonal covariance.

By assuming different priors for the invariant and spurious latent representations, inVAE can disentangle biological signals from technical noise. This disentanglement is particularly important for preserving the biological variability across datasets while mitigating the effects of technical artefacts.

### Loss function

The training of the inVAE model is conducted by optimising evidence lower bound (ELBO) that consists of a data reconstruction term and a regularisation term involving the KL divergence between the approximate posterior and the prior distribution. The ELBO is defined as:

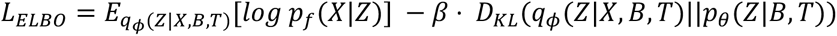

This term encourages the model to reconstruct the observed data accurately while regularising the latent space to follow the inferred prior distributions for *Z_I_* and *Z*_*S*_. In practice, we set β = 1 and perform a linear warm up for a small number of iterations from 0 to 1, such that first, a good reconstruction is ensured, and then the regularisation strength is increased. Disentanglement is implicitly encouraged as the number of latent dimensions for both the invariant and spurious latent space is restricted, and the reconstruction by the decoder is more effective if the information contained in the latent space is not duplicated. Furthermore, the conditionally independent priors on the latent spaces additionally encourage them to depend on the conditioning variables. We can further enforce independence between *Z_I_* and *Z*_*S*_ by minimizing their total correlation. The total correlation is a measure of dependence between two random variables, and penalizing this correlation encourages the model to learn statistically independent factors^72,73^:

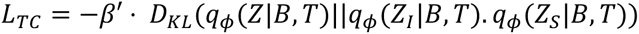

Measuring *q_ϕ_*(*Z*|*B*, *T*) requires the evaluation of the density 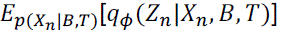, where *p*(*X_n_*|*B*, *T*) is the probability of observing sample *X_n_* ∈ *X* in domain (*B*, *T*). To solve this in practice, we extended the minibatch-weighted sampling proposed by Chen et al^73^.

### Data Representation

After training the model, we can use the learned encoder to represent the data as the mean of the underlying Gaussian distribution, using all available covariates, and can then optionally split the latent representation into the invariant and spurious parts.

### Prediction, reference mapping, and label transfer

For predicting annotations, e.g. cell types or diseases, a separate multilayer perceptron (MLP) can be trained using the invariant latent representation of the data. The MLP is trained in a supervised manner using the cell labels or metadata in the training data by predicting the labels from the invariant latent representation and backpropagating the cross-entropy loss of the predicted and ground-truth label.

Reference mapping is performed by employing the learned encoder and the available information in the form of gene counts and additional metadata and then taking the invariant part of the inferred mean of the latent distribution. In the straightforward approach, we simply use the known categories of the covariates or, for a new category, treat the corresponding one-hot encoded vector as all zeros. A more involved approach would infer the most suitable one-hot encoded vector for a new category. This can be done by selecting exactly one of the known categories that maximises the decoder density after passing all possible combinations to the encoder and decoder to reconstruct the input *X*. Another solution is to directly optimise the one-hot encoded vector by treating it as a parameter and performing backpropagation to maximise the decoder density with respect to the covariate vector. This results in a continuous version of the one-hot vector where multiple entries can be non-zero. In the classification experiments, we used the straightforward approach to show that our method already works well without using more complicated inference schemes.

After inferring the covariate vectors and the invariant latent representation, we can transfer labels by feeding this invariant latent presentation to the previously trained MLP to predict the reference labels.

### Hyperparameter

The inVAE model includes several key hyperparameters that impact both the training process and the quality of learned representations. In this section, we focus on hyperparameters specifically related to the model architecture, regularisation, and disentanglement objectives, along with default settings and alternative configurations.

Two of the most critical hyperparameters are the dimensions of the latent and spurious latent spaces, with default values of 30 and 5, respectively, for larger datasets. However, these values may be adjusted based on the number of cells, conditions, and the level of technical variation present in the data.

The neural network structure—shared across the encoder, decoder, and prior networks— includes several architectural parameters. These include the number of hidden layers, set by default to two; the hidden layer dimensions, set to 128; and the activation function, which is ReLU. Batch normalisation and dropout are also configurable, with default values set to True and a dropout rate of 0.1, respectively. We observed that these default values worked commonly well across all the datasets in this paper.

Conditional priors for latent variables can be modified to depend on selected covariates, as detailed in the following section. For the Gaussian distribution used in the priors, the default setting fixes the mean at zero while allowing the variance to be learned. However, depending on the data, the mean can be learned as well.

By default, the decoder models the observed data using a Negative Binomial distribution, although other distributions—such as Gaussian for normalised gene expression data or Poisson for raw count data—are also possible.

The batch size, set to 256, determines the number of samples processed per update of model parameters. Larger batch sizes generally yield more stable gradient estimates but require additional memory.

Lastly, an option is available to incorporate spurious covariates, such as batch ID, directly into the latent space via one-hot encoding, similar to the approach in scVI^19^. This setting bypasses the need to learn a separate spurious latent representation, effectively reducing the spurious latent dimension to zero. While this can help remove technical variation in some datasets, it should be used cautiously to avoid potential overcorrection.

### Covariates

Covariates can serve as inputs to either the conditional prior of the invariant latent space or the spurious latent space. The combined set of covariates is appended to the gene expression counts and then fed into the encoder of inVAE. Continuous covariates are input directly, with any necessary transformations applied beforehand, while categorical covariates are encoded as one-hot vectors, with the number of categories specified at initialization. For categorical covariates not observed during inference, all entries are set to zero. In this paper, all covariates are treated as categorical variables.

### Feature Importance

The feature importance of inVAE’s latent space is assessed by first training a separate random forest classifier from scikit-learn^44^ for each covariate, using the entire latent space (both invariant and spurious) across the dataset. Next, we used the ‘permutation_importanc’ function in scikit-learn^44^, with 10 repetitions, to estimate the importance of each latent variable. This approach permutes each feature individually and calculates the difference between the original score and the mean score of the permuted versions, providing a more robust estimate of feature importance, particularly for prediction targets with high cardinality, such as batch ID.

### Classification

Classification is conducted by initially training each integration method using its default or recommended hyperparameters, with the latent space dimension set to 30 for consistency across models. For cell type prediction, each method’s built-in classifier is utilised; in inVAE’s case, this is a shallow MLP with a single hidden layer of dimension 50. For the other prediction targets—disease status and combined cell type-disease status—a separate MLP with the same configuration is trained on each model’s latent space.

### Benchmarking of scRNA-seq Integration Methods

We benchmarked several tools to assess the performance of scRNA-seq integration (batch correction), including scVI^19^, Combat-PCA^36^, PCA, scPoli^26^, fastMNN^21^, scDisInFact^37^, scMerge-PCA^38^, scanorama^18^, and harmony^22^. The evaluation was conducted using the scib-metrics package^41^, calculating metrics based on embeddings generated by each method. For tools that output a corrected matrix rather than embeddings, we applied PCA to the corrected matrices for a standardised comparison. The input data was provided in h5ad format for direct access in Python, while R-based tools used data via the schard package^74^.

**Combat** was executed in Python using Scanpy’s combat function, specifying donor ID as the batch key. PCA was run with 25 dimensions for heart data and 30 for lung data, as Combat does not directly produce embeddings.

**scVI** was implemented using the scVI Python package, with donor ID set as the batch_key and n_latent dimensions set to 25 for heart and 30 for lung. Default parameters were used (n_layers=2, gene_likelihood=“nb”).

**Harmony** was run using a PyTorch-based version of the Harmony package (originally in R). First, we computed PCA embeddings with 25 components for heart data and 30 for lung data. These embeddings were then passed to the harmonise function, with donor ID specified as the batch_key.

**scMerge** was executed using the scMerge2 function from its R package, with donor ID set as the batch key and disease as the condition, employing default settings (k_celltype=20). Since scMerge does not output embeddings directly, PCA with 25 dimensions for heart and 30 for lung data was applied afterward.

**Scanorama** was applied using the scanorama package in Python. The input data was divided into separate objects based on donor ID (batch), and integration was performed using the integrate_scanpy function with 25 dimensions for heart and 30 for lung. The resulting embeddings from individual objects were then concatenated.

**fastMNN** was implemented using the batchelor package in R, with donor ID specified as the batch key and the number of dimensions (d) set to 25 for heart and 30 for lung. As fastMNN does not generate embeddings directly, we applied PCA with 25 dimensions for heart and 30 for lung to ensure consistency across methods.

## Metrics

### Data integration metrics

We evaluated the quality of data integration using established metrics for assessing single-cell data integration, all implemented in the scIB package^41^. Biological conservation was quantified using the following metrics: Cell Type Average Silhouette Width (ASW), Isolated Label F1, Isolated Label Silhouette, Normalised Mutual Information (NMI), and Adjusted Rand Index (ARI). Batch mixing was assessed with Graph Connectivity and Batch ASW.

**Normalised Mutual Information (NMI)** measures the overlap between two clusterings, calculated between cell type labels and clusters derived from unsupervised clustering of the integrated dataset. NMI values range from 0 to 1, where 1 indicates a perfect match between clusterings, and 0 indicates no overlap.

**Adjusted Rand Index (ARI)** assesses the similarity between two clusterings by considering both matching overlaps and correct disagreements. This score is calculated between the cell type labels and the clusters on the integrated dataset, ranging from 0 to 1, with 1 representing a perfect score.

**kBET (k-nearest neighbour Batch Effect Test)** evaluates whether the label composition within the k nearest neighbours of a cell matches the expected global label composition. This test is repeated across a random subset of cells, with results reported as a rejection rate across tested neighbourhoods, operating on a K-nearest-neighbour (KNN) graph.

**Cell Type Average Silhouette Width (ASW)** quantifies the relationship between the within-cluster distance of a cell and its between-cluster distance to the nearest other cluster. Scores range from –1 to 1, where –1 indicates complete misclassification, 0 indicates cluster overlap, and 1 indicates well-separated clusters. We use two versions of ASW: one based on cell type labels and a modified version for batch mixing (Batch ASW). The ASW score is rescaled to values between 0 and 1 using the following equation:

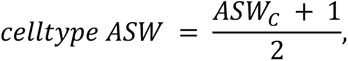

where C represents the set of all cell type labels.

**Batch ASW (Average Silhouette Width):** The batch ASW quantifies the quality of batch mixing in the integrated object. We obtain it by computing the ASW but on batch labels instead of cell type labels. Scores of 0 are indicative of good batch mixing, while any deviation from this score is the result of batch effects. In order to have a metric bound between 0 and 1, the following transformation is applied:

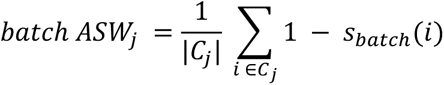

Here, *C_j_* is the set of cells with label *j* and |*C_j_*| is the support of the set.

The final score is obtained by averaging the batch ASW values obtained for each label:

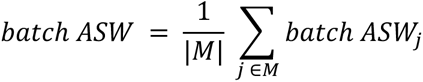

with *M* being the set of unique cell labels.

**Isolated label F1:** Isolated labels are defined as cell type labels which are found in the smallest number of batches. We aim to determine how well these cell types are separated from the rest in the integrated data. To do so, we find the cluster containing the highest number of cells from such isolated labels, and we then compute the F_1_ score of the isolated label against all other labels within the cluster. We use the standard F_1_ formulation:

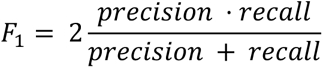

Once again, scores of 1 represent the desirable outcome in which all cells from the isolated labels are grouped in one cluster.

**Macro F1:** This score treats every class equally, irrespective of the class size, and is thus suitable for imbalanced datasets. It is calculated by an unweighted average of the F_1,*i*_ scores per class *i* as

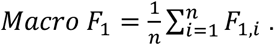

**Isolated label silhouette:** This is a cell type ASW score but computed only on the isolated labels.

**Graph connectivity:** This metric measures whether a KNN graph (G) computed on the integrated object connects cells that fall within the same cell type. It is bound between 0 and 1, with 0 indicating a graph where all cells are unconnected and, and 1 occurring when all cells of the same cell type are connected in the integrated output.

For each cell type label a subset graph G(N_c_, E_c_) is computed, the final score is obtained using the following formulation:

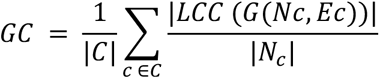

where *C* is the set of unique cell type labels, |*LCC* (*G*(*Nc*, *Ec*))| is the number of nodes in the largest connected component in the KNN graph and |*N*_*c*_| is the number of nodes in the graph.

### Gene module identification

We first applied inVAE to the entire spatial lung dataset^49^, followed by selecting AT1 and ciliated cells to examine spatial variation and identify related genes and pathways in specific locations.

For downstream analysis, we used the Hotspot (v1.1.1) software^50^ to identify gene modules in an unsupervised manner, leveraging the similarity graph (KNN) constructed from inVAE’s embedding. Hotspot performed a gene autocorrelation test on this KNN graph to detect genes exhibiting non-random spatial patterns. We then selected the top 1000 genes based on their autocorrelation scores for hierarchical clustering, which produced gene modules. These modules were visualised in a correlation plot accompanied by a dendrogram.

### Meta cells

We used two tools for inferring meta cells, each with slightly different features but both based on similarity graphs, such as KNN. SEACells^39^ (n_seacells=100) identified meta cells from inVAE’s embedding and assessed the purity of subpopulations, defined as the percentage of the most frequent cell type within each meta cell.

Milo^46^ (n_neighbors=10), was run on inVAE’s embedding and provided the mean developmental stage within each neighbourhood of cells.

### Pseudo ordering of cells

To derive pseudo order of the lung tip cells, we used the basic curved trajectory analysis workflow from scFates^47^ (v1.0.7), with default parameters as specified in the software’s tutorial. First, a principal curve was inferred, followed by the identification of root cells. Cells were then projected onto this principal graph to compute pseudo-orders. Genes significantly associated with cell state dynamics were identified and modelled as functions of their positions along the trajectory. These genes were then clustered based on their expression patterns. Finally, we visualised gene expression trends by plotting residuals—differences between observed and fitted expression values—against pseudo-orders.

### Pathway enrichment analysis

In the heart failure dataset, we ranked the differentially expressed genes (DEGs) using sc.tl.rank_genes_groups. We then selected the genes with adjusted p-value<0.05 and log fold change >0.0. We ran the EnrichR^75^ which is implemented in gseapy^76^ with Gene Ontology Database^77^ (gene_sets=’GO_Biological_Process_2023’) using the selected DEGs to find the enriched pathways. For the spatial lung datasets, we used the EnrichR with Wiki Pathways^78^ (gene_sets=’WikiPathways_2024_Human’) for the significant gene modules. The significant pathways are selected based on the threshold of adjusted p-value < 0.05.

### Gene score

We used scanpy.tl.score_genes to measure gene enrichment scores. This score is calculated as the average expression of a target gene set, minus the average expression of a reference gene set. The reference set is randomly sampled from the gene pool for each binned expression value.

## Data availability

All datasets used in this manuscript are publicly available and already published. The original papers regarding each dataset are cited in the manuscript. The processed data and train models are available at [link to the github].

## Code availability

The python package is available in [https://github.com/theislab/inVAE]. The preprocessing and downstream analysis pipelines are included in [https://github.com/theislab/inVAE_reproducibility].

## Contributions

H.A. conceptualised and designed the study. H.A. and F.K. developed the tool. F.K. wrote the package. H.A., K.B.M. and J.C. selected the datasets. H.A., F.K., D.P. and B.C. performed the analysis. H.A. wrote the first draft of the manuscript and made the figures. H.A., F.K., D.P. and B.C. wrote the Methods Section. H.A., D.P., K.B.M. and R.V. edited the manuscript. T.J., J.C., and S.A.T. proofread the heart results. D.P. and K.B.M. proofread the lung results. All the authors critically reviewed and approved the manuscript. R.V. and F.J.T funded the project.

## Acknowledgements

We would like to express our gratitude to Aidan Maartens for his insightful comments on the manuscript and A. García from Bio-Graphics for the illustration of Figure 1. We are also grateful to Ana Paredes and Kazumasa Kanemaru for fruitful discussions on heart data analysis and Peng He for his assistance with the fetal lung data. Additionally, we thank all members of Vento-Tormo, Theis, and Teichmann laboratories for their valuable discussions and feedback throughout this project. We also extend our thanks to everyone who has used our tool, providing invaluable input that helped us improve inVAE. We are especially grateful to the CellGen admin team at the Wellcome Sanger Institute for their continuous support with computational resources and infrastructure. We acknowledge that Figures 2a, 4a-5a are depicted using bioRender.

## Disclosure of Potential Competing Interest

F.J.T. consults for Immunai Inc., CytoReason Ltd, Cellarity, and BioTuring Inc. and has an ownership interest in Dermagnostix GmbH and Cellarity. In the past 3 years, S.A.T. has consulted for or been a member of scientific advisory boards at Qiagen, Sanofi, Element Biosciences, Xaira Therapeutics, GlaxoSmithKline and ForeSite Labs. She is a co-founder and an equity holder of TransitionBio and EnsoCell. She is a part-time employee at GlaxoSmithKline and a non-executive director of 10x Genomics.

## Supplementary Figures and Notes

**Supplementary Figure 1:**
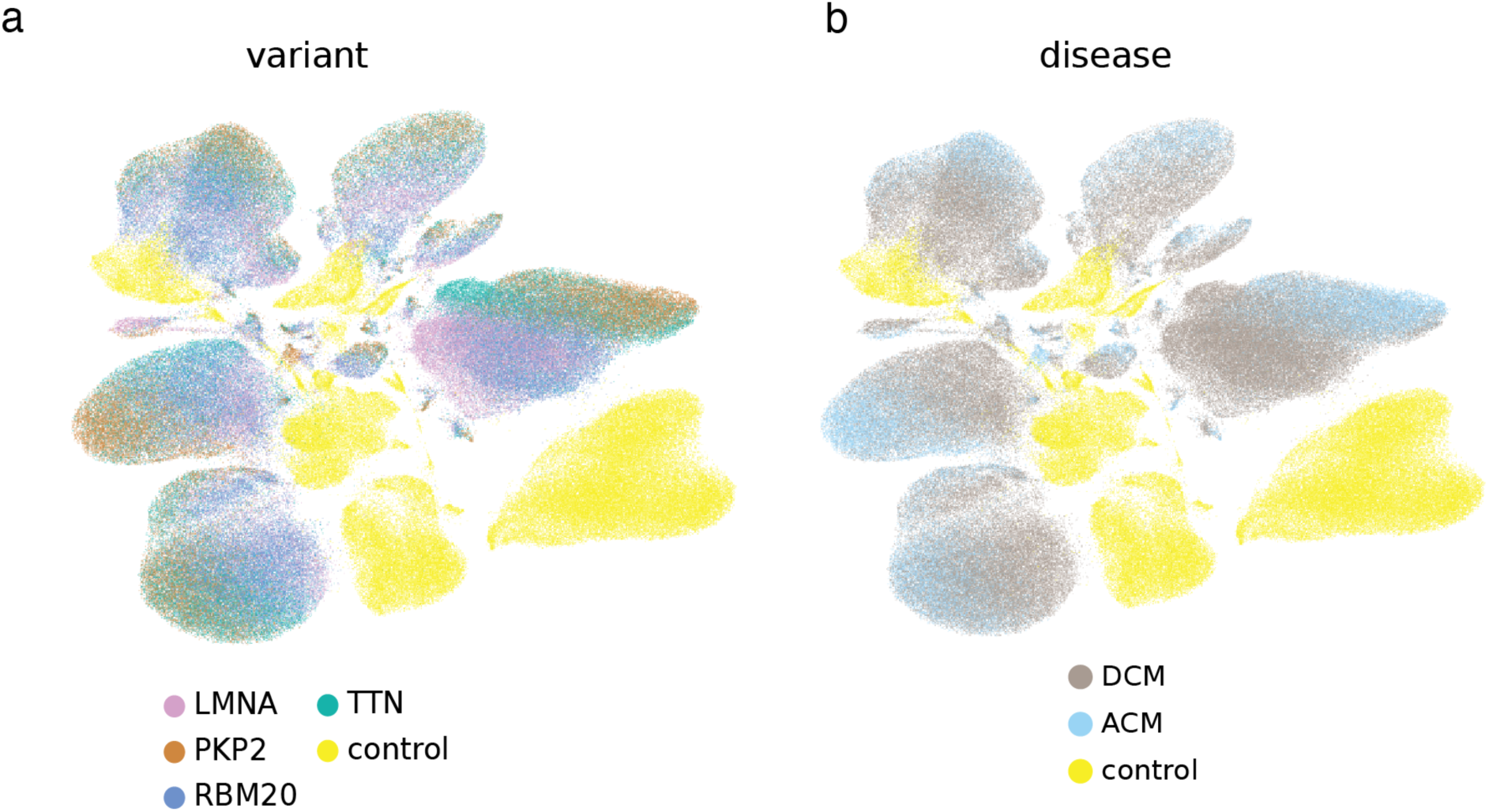
Representation of heart atlas (related to Figure 2). **a**, The UMAP visualisation of the heart atlas coloured by variants, and **b,** diseases.

**Supplementary Figure 2:**
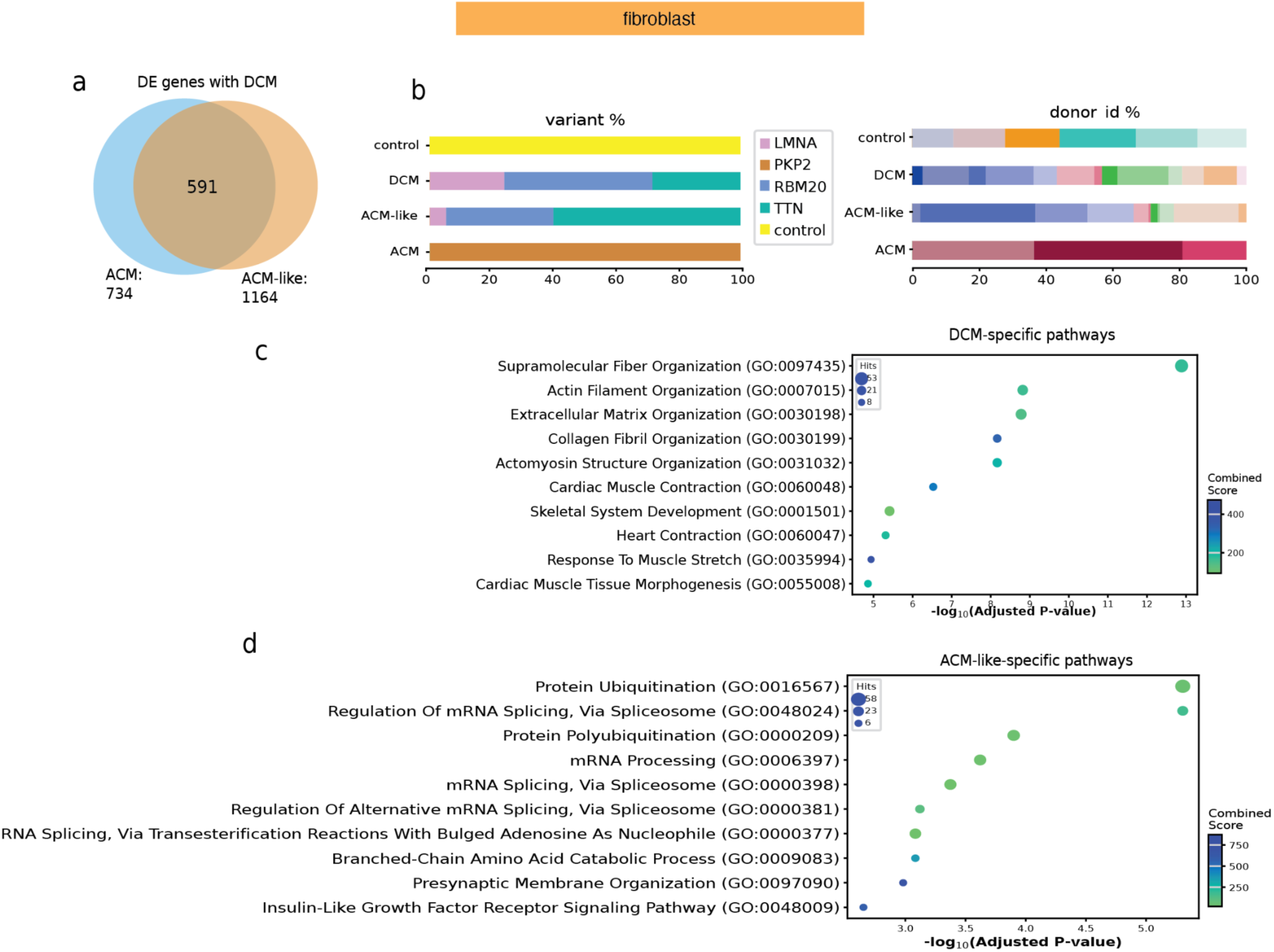
Downstream analysis of fibroblast representation in the human heart atlas (related to Figure 2). **a**, Shared differentially expressed genes between ACM and ACM-like cells compared to DCM cells are shown. **b,** The compositional analysis of diseased and control fibroblast populations is presented. **(c, d),** Enriched pathways in DCM and ACM-like cells are identified using GSEApy^76^. ACM, arrhythmogenic cardiomyocyte; DCM, dilated cardiomyocyte.

**Supplementary Figure 3:**
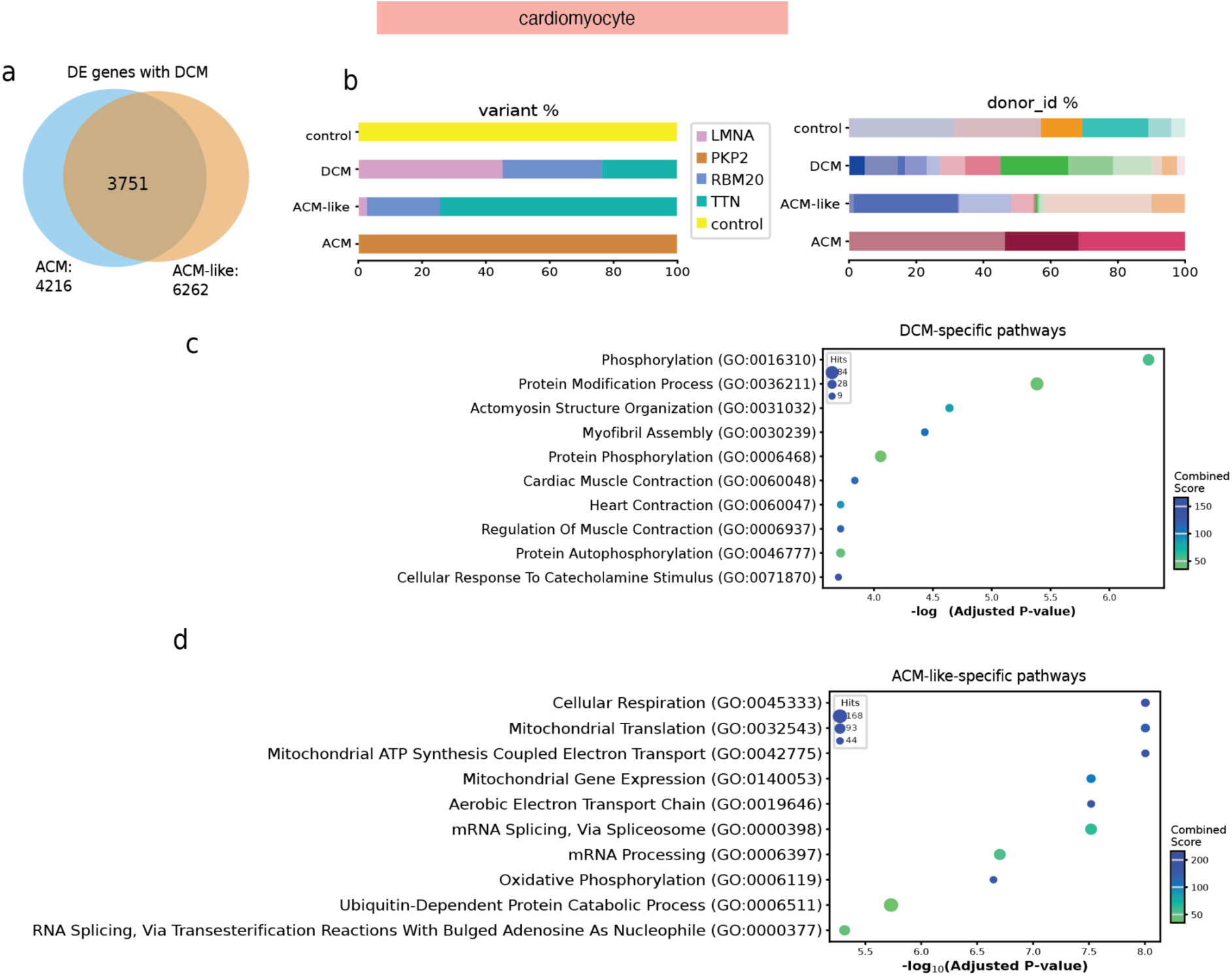
Downstream analysis of cardiomyocyte representation in the human heart atlas (related to Figure 2). **a**, Shared differentially expressed genes between ACM and ACM-like cells compared to DCM cells. **b,** Compositional analysis of diseased and control cardiomyocyte populations. **(c, d),** Enriched pathways in DCM and ACM-like cells are identified using GSEApy^76^. ACM, arrhythmogenic cardiomyopathy; DCM, dilated cardiomyopathy.

**Supplementary Figure 4:**
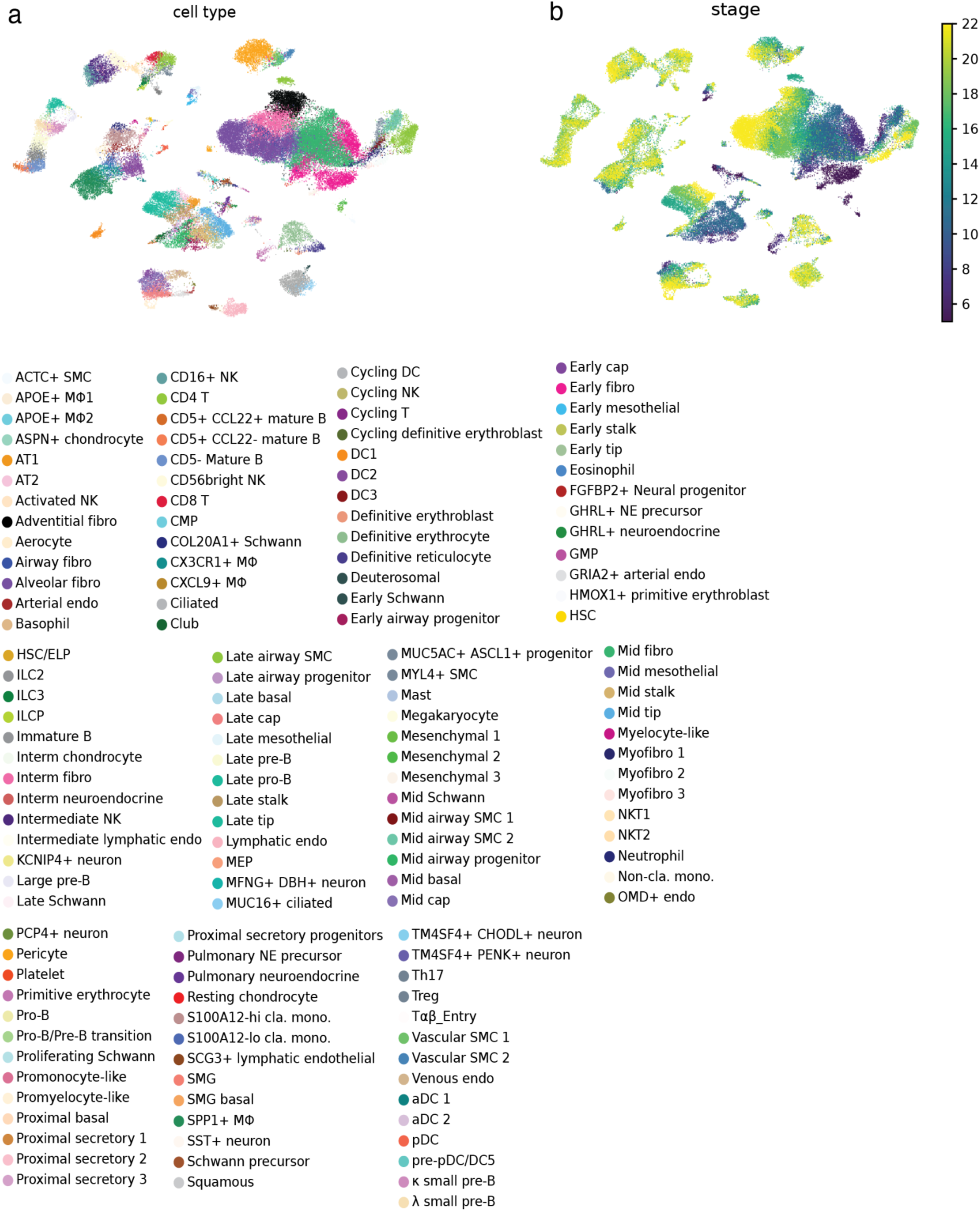
Representation of fetal lung atlas (related to Figure 4). **a**, Uniform Manifold Approximation and Projection (UMAP) visualisation of the fetal lung atlas coloured by cell types, and **b,** developmental stages.

**Supplementary Figure 5:**
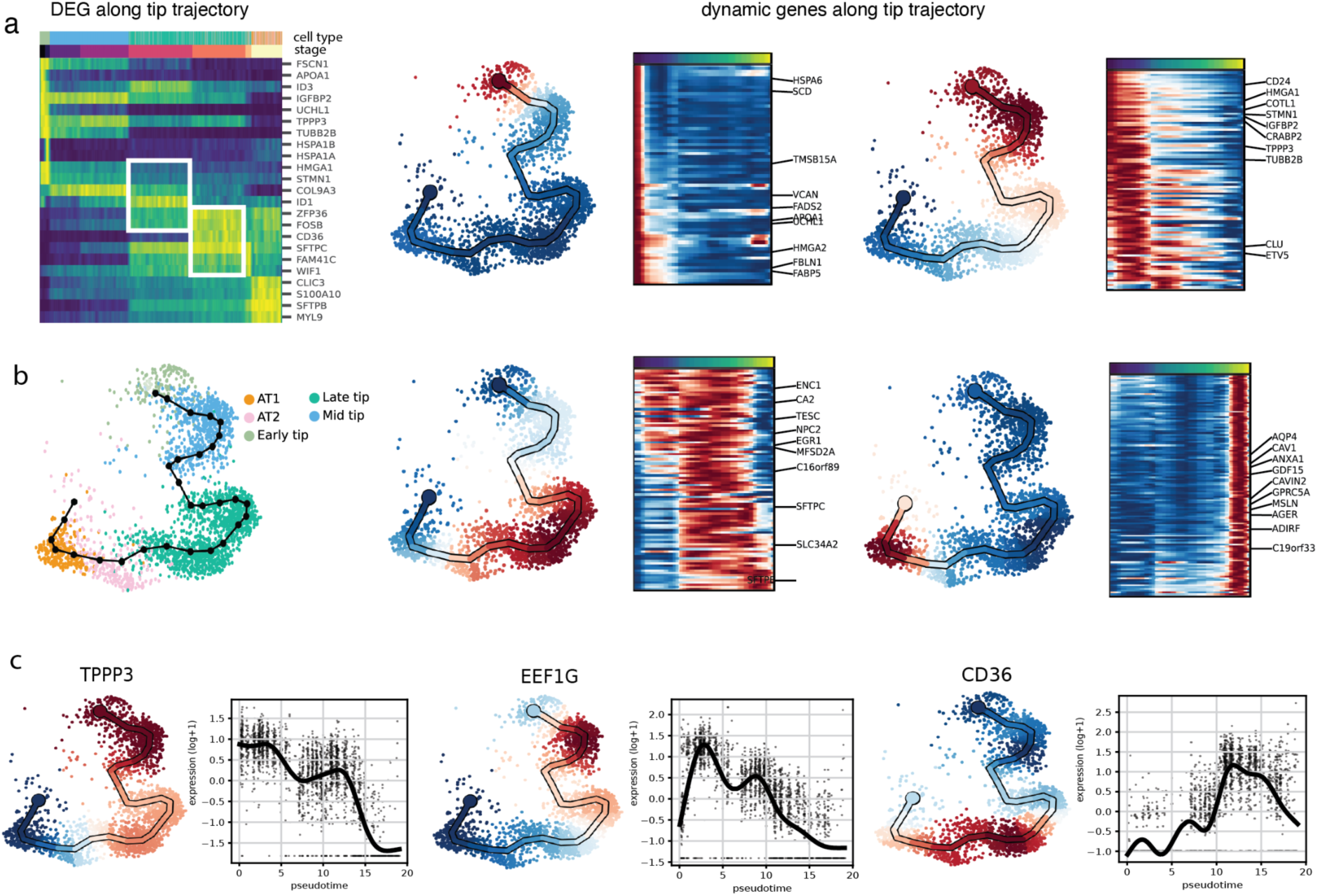
Downstream analysis of tip trajectory (related to Figure 4). **a**, Differentially expressed genes along the tip temporal states identified using scanpy.rank_genes. **b,** The pseudo-ordering of cells and dynamically expressed genes along the trajectory, analysed with scFates^47^. Each plot shows gene modules differentially expressed in different parts of the manifold. **c,** Expression patterns of three selected differentially expressed genes along the trajectory.

**Supplementary Figure 6:**
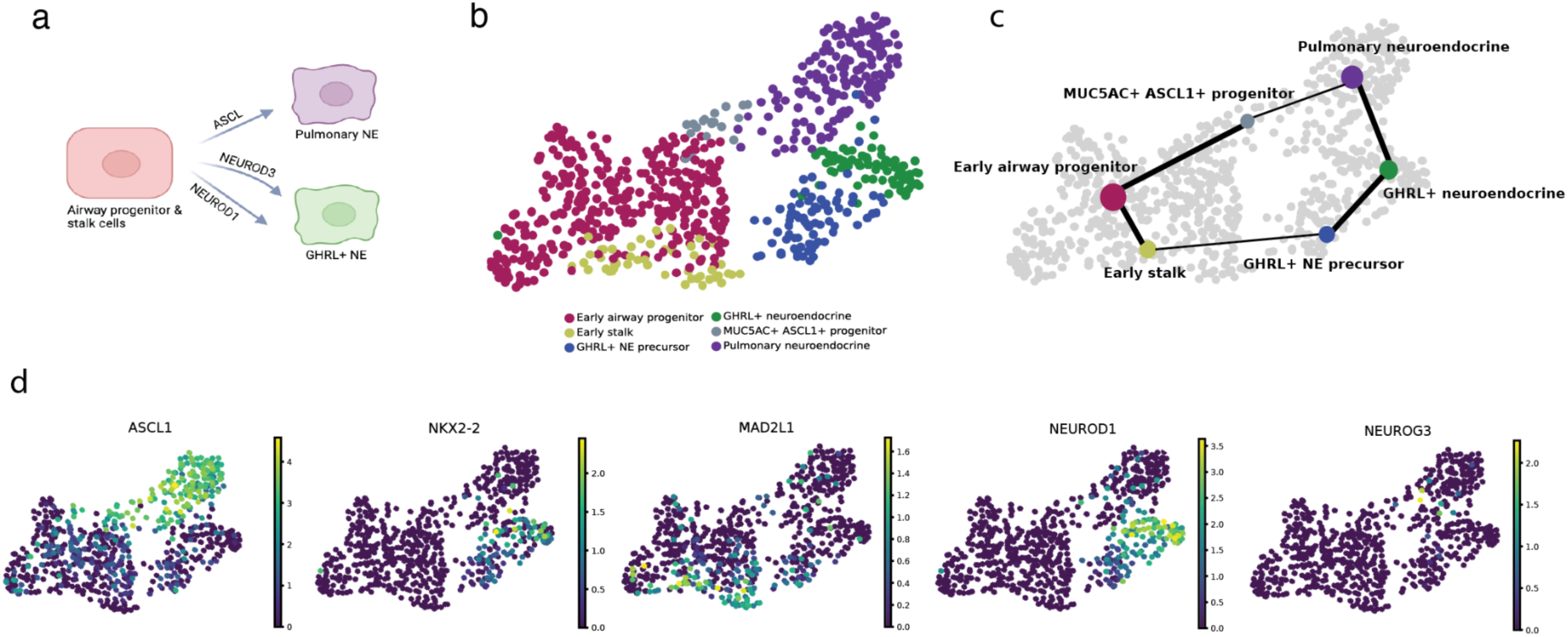
Downstream analysis of neuroendocrine trajectory (related to Figure 4) **a**, The neuroendocrine differentiation process described by He, et al.^45^. **b,** Corresponding cells from the atlas are coloured by cell type. **c,** Trajectory analysis using PAGA depicts the differentiation trajectories. **d,** The expression profiles of key neuroendocrine biomarkers, including *ASCL1*, *NKX2-2*, *MAD2L1*, *NEUROD1*, and *NEUROG3*, are shown along the trajectory.

**Supplementary Figure 7:**
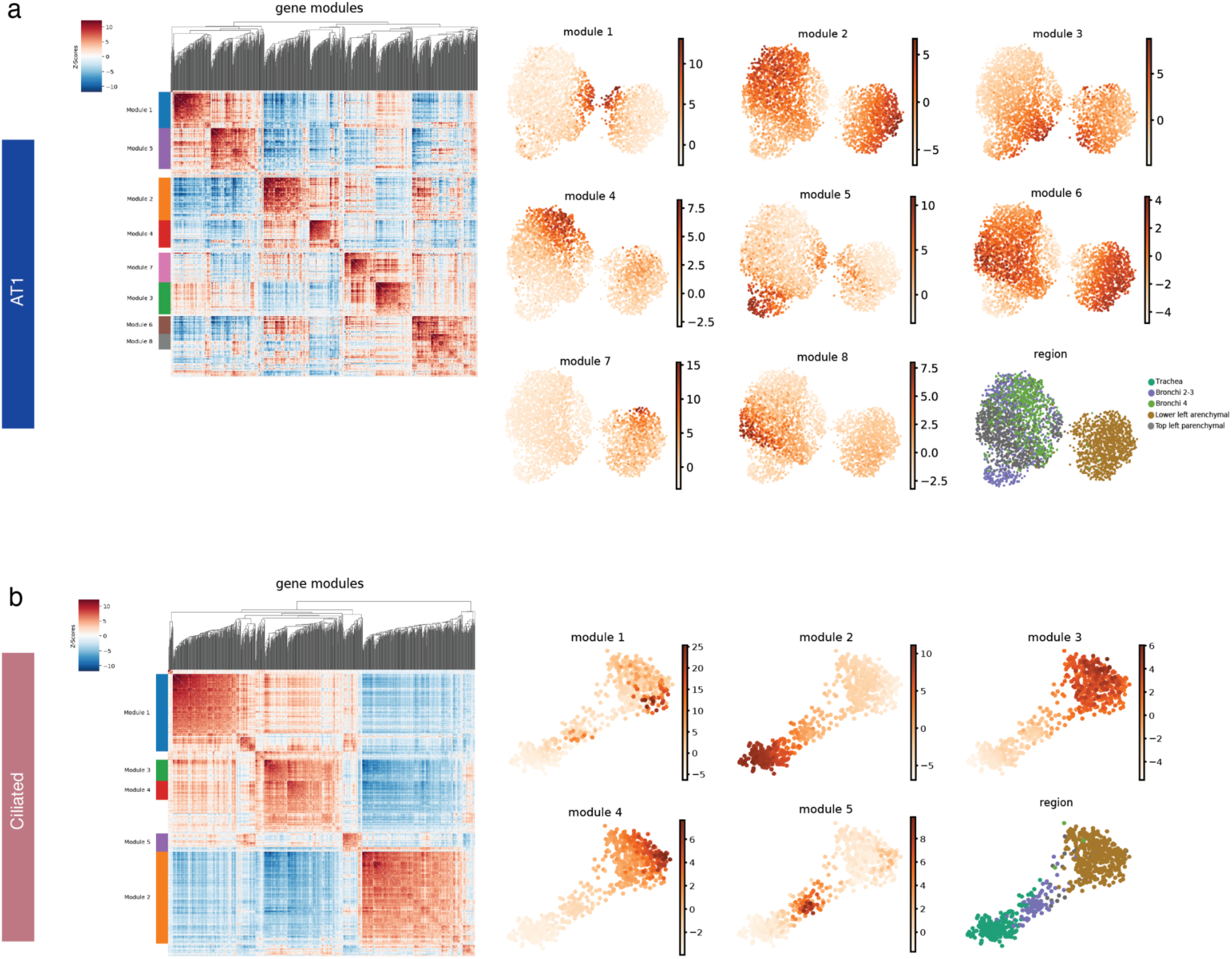
Spatially expressed gene modules in human lung (related to Figure 5) **a**, Gene modules that are differentially expressed in different neighbourhoods on the inVAE embedding for AT1 cells using Hotspot^50^. The heatmap shows the correlation between the genes followed by their corresponding gene score and their associations with different spatial locations of the lung. **b,** Gene modules for ciliated cells and their corresponding score.

**Supplementary Figure 8:**
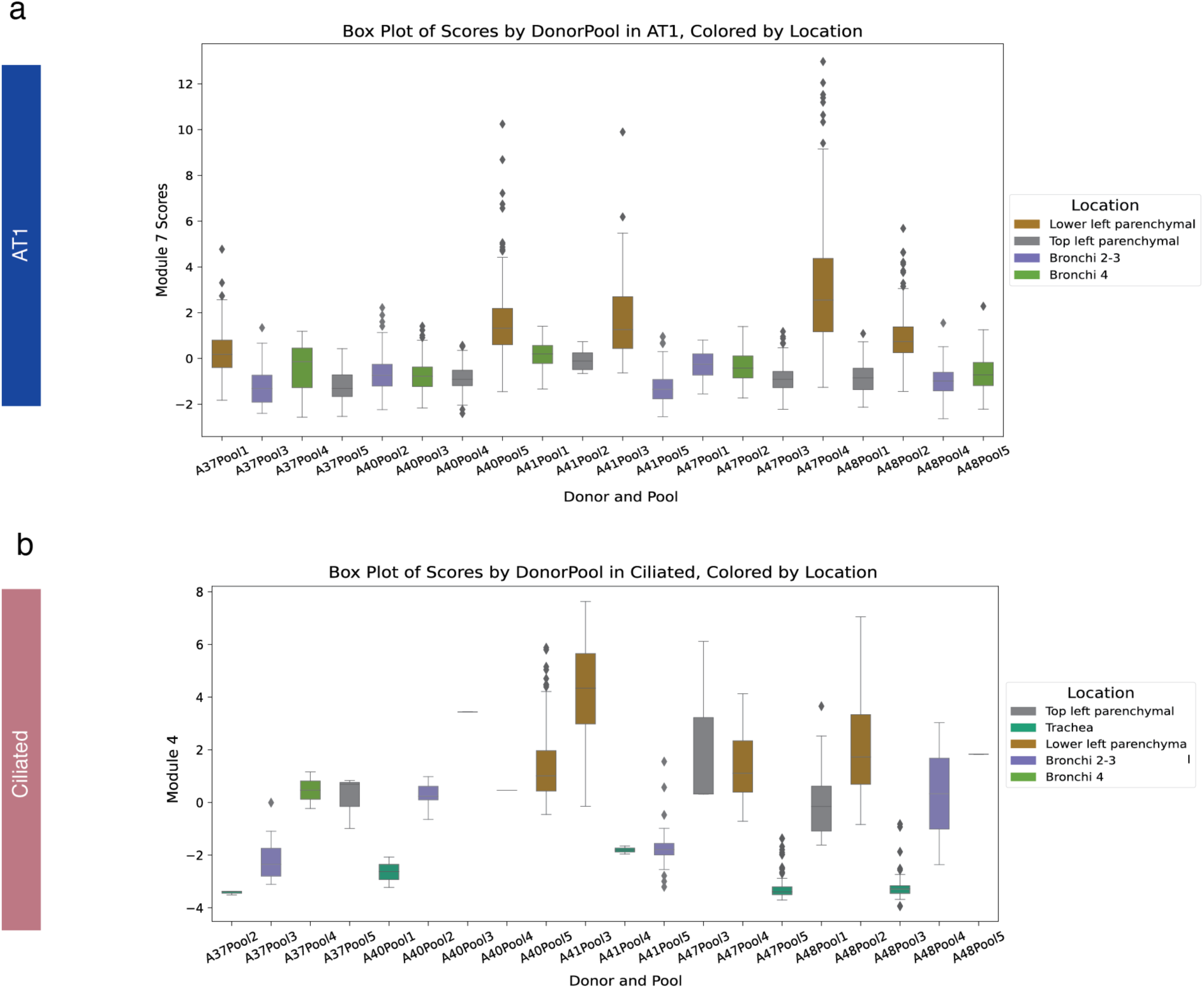
Cell type-specific gene module score distribution across donors and pools (related to Figure 5) **a**, Distribution of module 7 scores in alveolar type 1 (AT1) cells stratified by donors and pools. Donors and pools are mapped across distinct anatomical regions of the lung as indicated in the legend. **b,** Module 4 score distribution in the Ciliated cells across donors and pools.

## Notes

### Competing Interest Statement

The authors have declared no competing interest.

